# Screening yeast display libraries against magnetized yeast cell targets enables efficient isolation of membrane protein binders

**DOI:** 10.1101/728436

**Authors:** Kaitlyn Bacon, Matthew Burroughs, Abigail Blain, Stefano Menegatti, Balaji M. Rao

## Abstract

When isolating binders from yeast displayed combinatorial libraries, a soluble, recombinantly expressed form of the target protein is typically utilized. As an alternative, we describe the use of target proteins displayed as surface fusions on magnetized yeast cells. In our strategy, the target protein is co-expressed on the yeast surface with an iron oxide binding protein; incubation of these yeast cells with iron oxide nanoparticles results in their magnetization. Subsequently, binder cells that interact with the magnetized target cells can be isolated using a magnet. Using a known binder-target pair with modest binding affinity (K_D_ ~ 400 nM), we showed that a binder present at low frequency (1 in 10^5^) could be enriched >100-fold, in a single round of screening, suggesting feasibility of screening combinatorial libraries. Subsequently, we screened yeast display libraries of Sso7d and nanobody variants against yeast displayed targets to isolate binders specific to the cytosolic domain of the mitochondrial membrane protein TOM22 (K_D_ ~ 271-2009 nM) and the extracellular domain of the c-Kit receptor (K_D_ ~ 131 to K_D_ >2000 nM). Additional studies showed that the TOM22 binders identified using this approach could be used for the enrichment of mitochondria from cell lysates, thereby confirming binding to the native mitochondrial protein. The ease of expressing a membrane protein or a domain thereof as a yeast cell surface fusion – in contrast to recombinant soluble expression – makes the use of yeast-displayed targets particularly attractive. Therefore, we expect the use of magnetized yeast cell targets will enable efficient isolation of binders to membrane proteins.

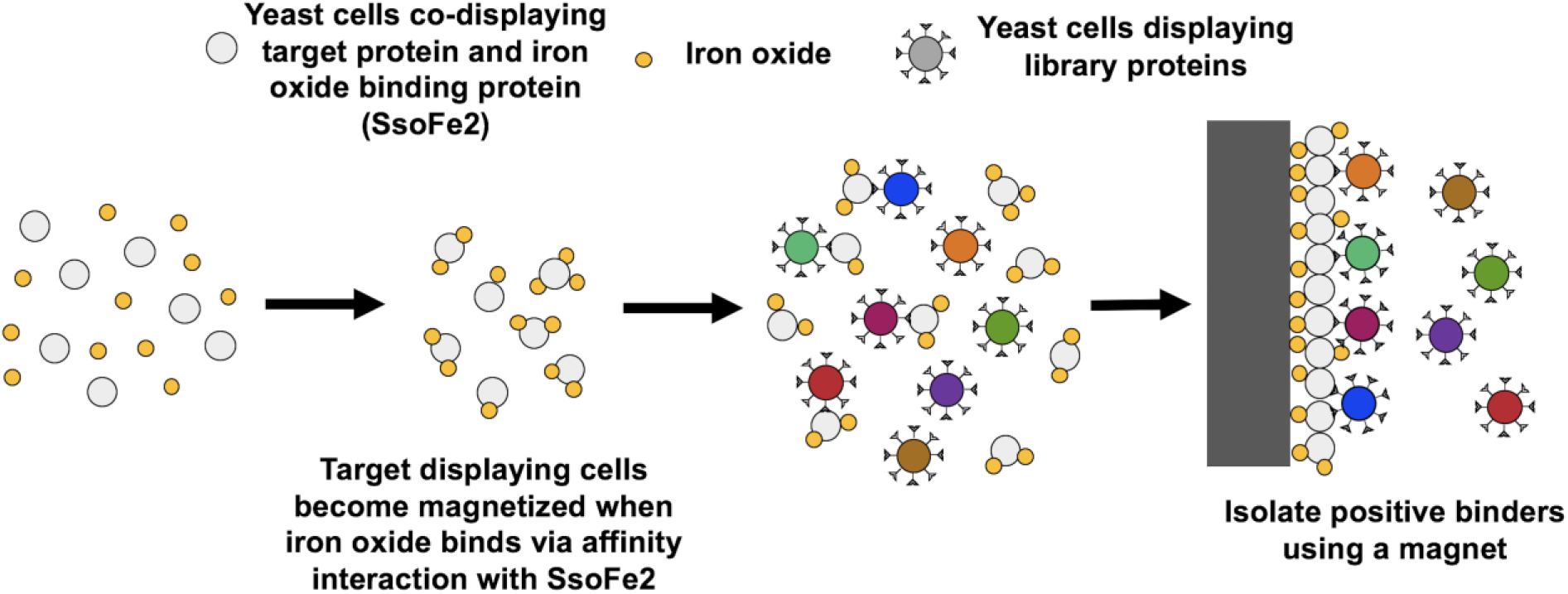

## Introduction

Binding proteins that specifically target membrane proteins are important in several applications, including therapeutics, diagnostics, and the separation of biologics such as organelles and cells. Disease states are often defined by the overexpression of one or more specific membrane proteins^1,2^. Therefore, binding proteins with specificity to these targets are commonly used to direct drug therapies^3,4^, to monitor disease advancement using imaging probes^5,6^, or to predict disease progression^2^. Furthermore, advances in proteomics and genomics are enabling the identification of new clinically relevant membrane protein targets^7,8^. Binders to membrane proteins are also used to isolate intracellular organelles for basic research or to separate cell populations for transplant purposes^9–11^.

Several high throughput combinatorial library screening platforms, such as phage display^12,13^, yeast display^14^, or mammalian display^15,16^, have been used to identify ligands with unique or enhanced binding affinity for a target of interest. Nevertheless, isolation of binders to membrane protein targets can be challenging. Strategies for the isolation of binders to membrane proteins typically utilize a recombinantly expressed extracellular domain of a target protein. Recombinantly produced protein is often misfolded or adopts a conformation that is different from the native protein^17,18^. Misfolding is also compounded by storage and purification conditions that can promote aggregation^19^. Additionally, recombinant proteins are often fused with biological or chemical tags to assist in purification or selection. Binders with affinity for these tags rather than the protein may be isolated^14,20^. Consequently, use of recombinant protein targets may result in the isolation of binders that interact with epitopes that are inaccessible or not present when the membrane protein is natively expressed^21^. These challenges may be overcome by screening libraries against whole cell targets expressing the membrane protein^22^. However, here too, isolation of specific binders may be difficult as the expression levels of the target protein may be low while other cell surface proteins are present at high density^23^.

As an alternative to the aforementioned approaches, yeast surface display has emerged as an attractive strategy for expressing membrane protein targets in the context of structure-function studies as well as screening phage display libraries^24–30^. In yeast surface display, a protein of interest is expressed as an N- or C-terminal fusion to the Aga2 subunit of the yeast mating protein a-agglutinin^14,31^. The Aga2 subunit in turn is linked to the cell wall associated Aga1 subunit, tethering the protein of interest to the cell wall. Notably, Zhao *et al.* have used yeast surface display to express a variety of extracellular membrane domains^25^. Here, we describe the development and evaluation of a screening strategy wherein yeast displayed membrane protein targets are used for the isolation of binding proteins from a yeast display library.

The ease of expressing a membrane protein or a domain thereof as a cell surface fusion – in contrast to recombinant soluble expression – makes the use of yeast-displayed targets particularly attractive^32^. However, the critical challenge in screening yeast display libraries using yeast displayed targets is the separation of binder cells from the non-binders. To efficiently enable this separation, we investigated a strategy wherein an iron oxide binding protein (SsoFe2) is used for the magnetization of yeast cells displaying a target protein (**Figure 1**)^33^. Briefly, the incubation of yeast cells co-expressing a target protein and SsoFe2 with iron oxide nanopowder results in the magnetization of target displaying yeast. Subsequently, the magnetized target yeast cells are incubated with a yeast display library, and any putative binder cells that complex with the target-displaying yeast are separated using a magnet.

**Figure 1.**
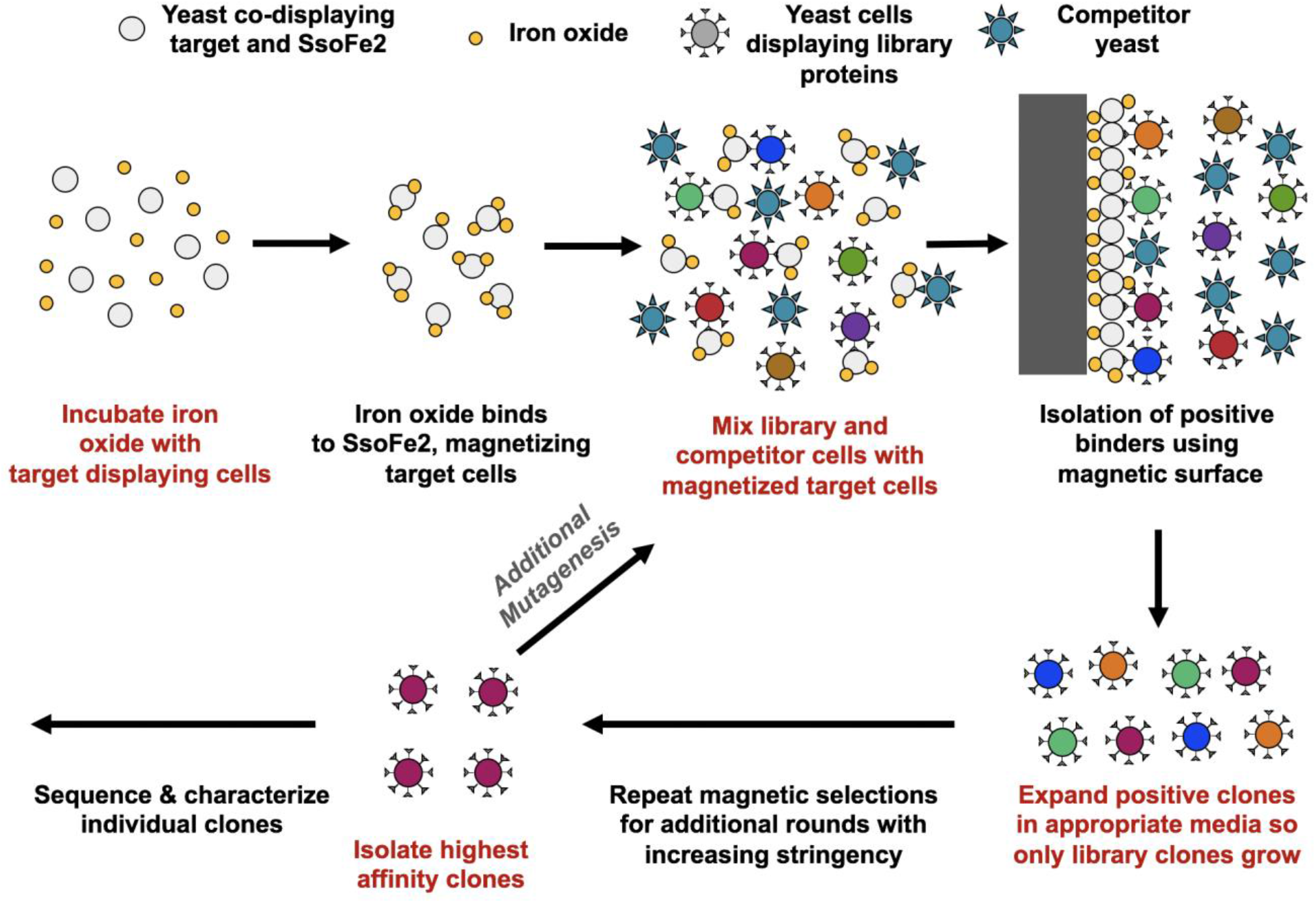
Overall strategy for screening a yeast displayed combinatorial library against yeast cells displaying a target protein. The target cells co-express the target protein along with SsoFe2, a protein with affinity for iron oxide. Incubation of the target cells with iron oxide nanoparticles enables magnetization of the target cells. After, the yeast display library is incubated with the magnetized target cells. Competitor yeast cells (cells displaying an irrelevant protein or no protein) are included to reduce non-specific binding. Any library cells that bind the target displaying cells can be expanded in appropriate media and screened further with increasing stringency to isolate higher affinity binders. Subsequently, individual clones can be sequenced and characterized.

We first demonstrated the feasibility of our strategy by performing quantitative studies focused on the enrichment of yeast cells displaying a model binder protein from a mixed population containing non-binder cells. Here, cells displaying the binder were recovered using magnetized yeast displaying the corresponding target protein. Subsequently, we evaluated our screening strategy for the isolation of binding proteins to two membrane protein targets of interest (TOM22 and c-Kit) from two yeast display libraries based on the Sso7d and nanobody scaffolds. Binding proteins for a wide spectrum of targets have been previously isolated from combinatorial libraries derived from these scaffolds^34,35^. TOM22 is a membrane-associated mitochondrial protein; c-Kit is a cell surface receptor protein that is expressed on hematopoietic stem cells. Notably, antibody binders to the cytosolic domain of TOM22 are used in commercially available reagent kits for the isolation of mitochondria from cell lysates; these reagents serve as a benchmark when evaluating the binding proteins isolated using our strategy.

## Results and Discussion

### Co-expression of two proteins as yeast cell surface fusions

To obtain selective expansion of the binder cells that complex with the magnetized target cells in culture, the target-displaying yeast cells and the yeast displayed library cells must utilize yeast surface display plasmids with distinct nutritional selection markers. Towards that end, we sought to co-express the target protein and the iron-oxide binding protein, SsoFe2, as Aga2 fusions using a single plasmid. Two different approaches were investigated, involving the use of a T2A ribosomal skipping peptide from *Thosea asigna* (**Figure 2A**) or a bidirectional *GAL1/GAL10* promoter (**Figure 2B**). The T2A sequence has been successfully used to translate multiple proteins from a single mRNA transcript in *S. cerevisiae*^36,37^. The native yeast display construct contains the bidirectional *GAL1/GAL10* promoter^38^.

**Figure 2.**
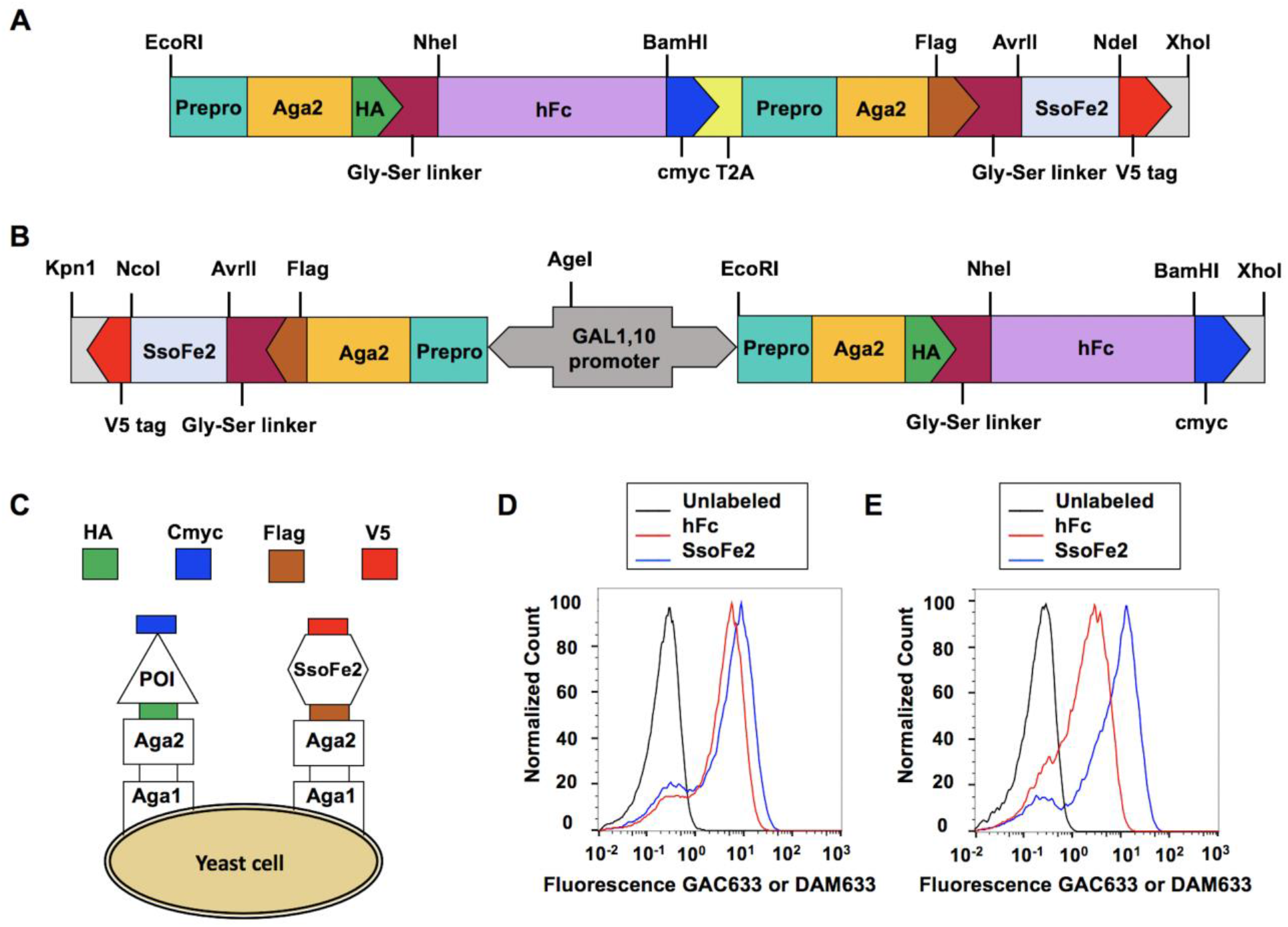
Constructs designed for the simultaneous display of two proteins on the surface of yeast as fusions to Aga2. In this example, the Fc portion of human IgG (hFc) and SsoFe2 are co-displayed as cell surface fusions. (**A**) Plasmid construct to achieve dual display of two proteins from a single mRNA transcript using a T2A ribosomal skipping peptide. (**B**) Plasmid construct to achieve dual display of two proteins using a bidirectional *GAL1/GAL10* promoter. In this example, hFc is under the direction of the *GAL1* promoter while SsoFe2 is under the direction of the *GAL10* promoter. (**C**) Visualization of protein co-expression as fusions to yeast cell surface including expression tags. (**D**) Representative flow cytometry analysis of hFc (blue) and SsoFe2 (red) expression by anti-cmyc and anti-V5 labeling, respectively, compared to unlabeled cells (black) using the ribosomal skipping construct. (**E**) Representative flow cytometry analysis of hFc (blue) and SsoFe2 (red) expression by anti-cmyc and anti-V5 labeling, respectively, compared to unlabeled cells (black) using the bidirectional promoter construct.

The functionality of the two co-expression approaches was evaluated for the simultaneous yeast surface display of the Fc portion of human IgG (hFc) and SsoFe2 (**Figure 2C**). For each construct, the expression levels of hFc and SsoFe2 were evaluated by immunofluorescent detection of the c-myc and V5 epitope tags, respectively, using flow cytometry analysis. Both approaches resulted in the co-expression of hFc and SsoFe2 as cell surface fusions (**Figures 2D, E**). However, the cell surface expression level of SsoFe2 was higher when the T2A peptide was used, relative to the case when SsoFe2 expression was under control of the *GAL10* promoter using the bidirectional promoter construct. Our results are consistent with previous studies by Rosowki *et al*^37^. The level of protein expression has been shown to be up to five-fold higher when the *GAL1* promoter was used instead of *GAL10*^39,40^. However, display levels are protein specific^41^; use of a stronger promoter may be desirable for proteins that are poorly displayed on the yeast cell surface relative to SsoFe2.

We chose to use the plasmid based on ribosomal skipping to achieve dual display in subsequent experiments because the cell surface display level was similar for the two co-expressed proteins during the initial evaluation of this construct. Note, however, the mechanism of ribosomal skipping and its effect on protein display must be considered when choosing a strategy for the co-display of a target protein and SsoFe2. 2A ribosomal skipping peptides share a conserved GDVEXNPGP sequence. Ribosomal skipping results in “cleavage” due to a failure to form a peptide bond between the C-terminal glycine and proline of this sequence during translation^42,43^. In the context of our plasmid construct, a major portion of the 2A peptide is fused to the C-terminus of the c-myc tag associated with the first translated protein (the target protein). It is important to note that the presence of these additional residues may affect the folding of certain proteins. More importantly, residues in the target protein upstream of the 2A sequence can affect the efficiency of ribosomal skipping. Finally, a proline residue is added to the prepro sequence associated with the second protein (SsoFe2) upon T2A cleavage; however, this did not affect surface display of SsoFe2.

### Yeast cells displaying a known binder protein can be enriched using magnetized yeast cells displaying the binder protein’s interaction partner

We conducted enrichment studies on a known binder-target pair to quantitatively assess the likelihood of isolating binders from a combinatorial library using magnetized yeast displaying the target protein. Yeast cells co-expressing hFc and SsoFe2 as cell surface fusions were magnetized by incubation with iron oxide powder (**Figure 3A**). We investigated the enrichment of cells expressing a previously identified Sso7d mutant (Sso7d-hFc) that binds hFc with modest binding affinity (K_D_~ 400 nM)^44^. Cells displaying Sso7d-hFc or the cytosolic domain of TOM22, an irrelevant protein, were mixed in varying ratios (10-100,000x excess of the cells displaying TOM22). The Sso7d-hFc/TOM22 cell mixtures were incubated with magnetized yeast cells expressing hFc as the target. The TOM22 cells act as competitors to the Sso7d-hFc cells for interaction with the target hFc cells. Subsequently, the iron oxide and any complexed cells were separated with a magnet. The number of Sso7d-hFc and TOM22 cells recovered by the magnetic target cells was quantified by plating on selective agar plates and colony counting (**Figure S1, Supplementary Methods and Discussion**).

**Figure 3.**
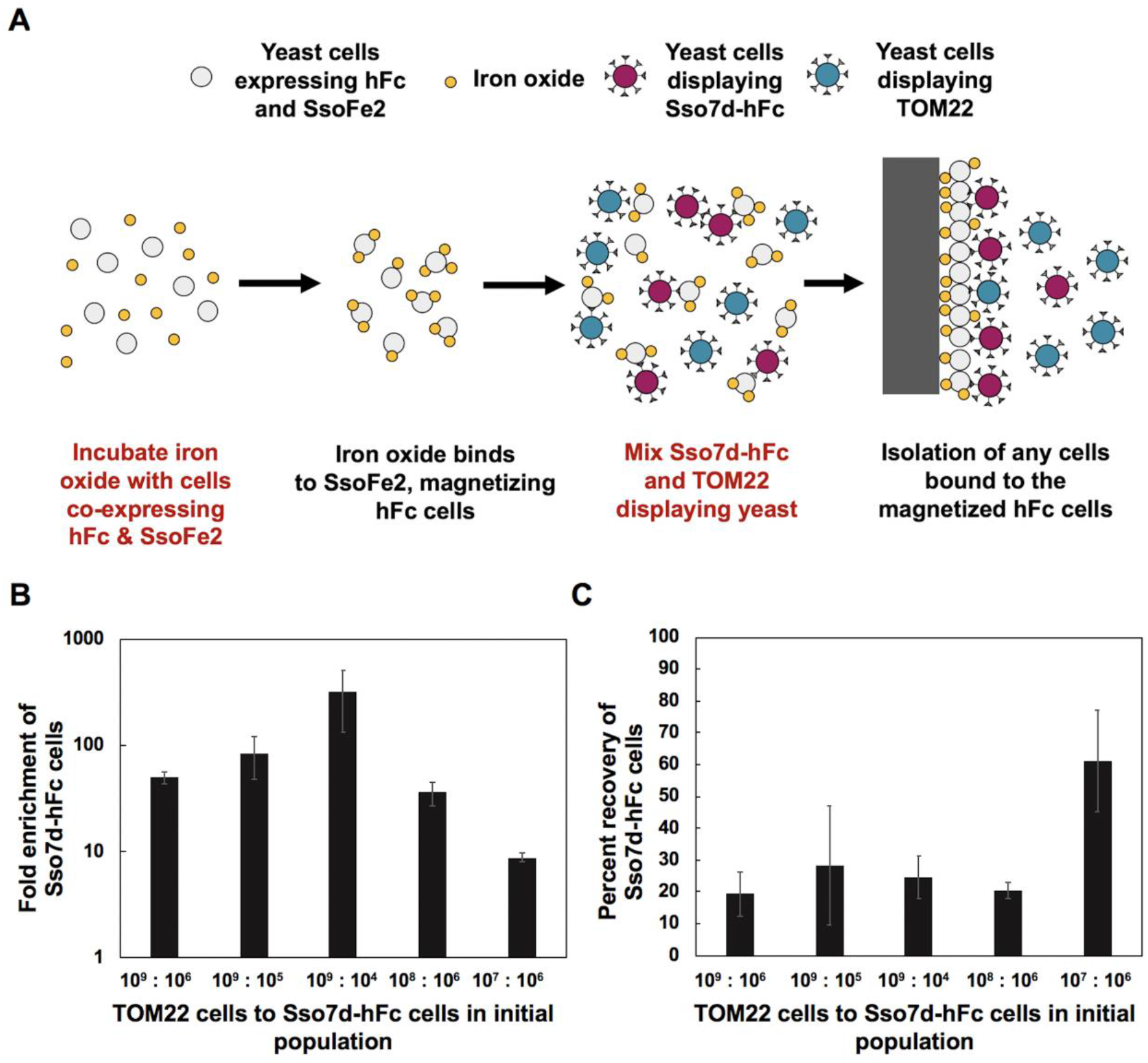
(**A**) Isolation of Sso7d-hFc displaying yeast using magnetic yeast co-expressing hFc and SsoFe2 in the presence of competitor yeast cells displaying TOM22. The hFc expressing cells are magnetized via iron oxide binding to SsoFe2 that is co-expressed on the cell surface. The number of TOM22 and Sso7d-hFc yeast cells was altered to vary the extent of competition from the TOM22 cells. Cells bound to the magnetic hFc cells were plated and quantitatively assessed to characterize the enrichment and recovery of the Sso7d-hFc population as well as the competitor TOM22 population. (**B**) Enrichment of Sso7d-hFc displaying yeast comparing the percentage of Sso7d-hFc cells in the recovered population to the percentage of Sso7d-hFc cells in the initial population. (**C**) Percentage of Sso7d-hFc displaying yeast in initial population that were recovered by the magnetic target cells. Error bars correspond to the standard error of the mean from three independent replicates.

The Sso7d-hFc displaying cells were enriched for all population ratios tested (**Figure 3B**), and at least ~20% recovery of these cells was observed (**Figure 3C**). Notably over 100-fold enrichment at ~ 20% recovery was observed when TOM22-displaying cells were present at 10^5^-fold excess. Based on the observed ~ 10-100-fold enrichment of binders in one round, one can reasonably assume that 4-5 rounds of screening with magnetized yeast cell targets would suffice, in general, to isolate specific binders from yeast display libraries. Additionally, a recovery of ~ 20% suggests that the library should be oversampled by at least five-fold, when screening using magnetized cell targets.

It is instructive to compare our results with those reported by Stern *et al.* wherein they assessed the recovery of binder yeast upon panning against adherent cells expressing the target^45^. In that study, binder cells (K_D_ ~ 2 nM) were mixed with non-binders at 1000x excess and recovered at ~ 100-fold enrichment and ~ 35% recovery; these values are comparable with our results. Interestingly, however, yeast cells displaying a low affinity binder (K_D_>600 nM) required multiple rounds of screening for enrichment^45^. In contrast, our approach could enrich a binder with K_D_ ~ 400 nM in a single round. Apart from the slightly higher binding affinity of Sso7d-hFc, the higher enrichment observed using our strategy may be attributed to there being a greater number of target cells present. We estimate that our screens contain at least 10x more target cells than when adherent cells on tissue culture plates are used as the target. The simplicity of increasing the number of magnetized target cells is a major advantage of our approach over classical cell panning using adherent cells as the scale up for large library screens is more efficient and even weak affinity binders can be isolated.

It is also important to note that yeast cells can be magnetized without the display of an iron oxide binding protein by simply incubating yeast cells with iron oxide nanopowder due to electrostatic interactions between the iron oxide and yeast cell surface proteins^33^. Therefore, non-specific adsorption of iron oxide to the yeast cell surface is an alternative strategy for obtaining magnetized yeast cell targets. However, use of SsoFe2 enables more robust cell magnetization. Wash steps and extended incubation in buffers containing high concentrations of carrier protein (e.g. bovine serum albumin (BSA)) can result in the loss of magnetization, presumably due to the dissociation of iron oxide particles from the yeast surface. The electrostatic interactions between the yeast cell surface proteins and the iron oxide are too weak to retain the iron oxide on the yeast surface when faced with competition from carrier protein or non-magnetized cells. Loss of magnetization has two deleterious consequences when screening combinatorial libraries – library binder cells associated with the de-magnetized target cells will be lost during the magnetic selection step, and non-specific re-binding of iron oxide to non-binder cells may result in their unwanted isolation. Our studies show that the loss of magnetization is significantly lower when SsoFe2 is present on the yeast cell surface (**Figure S2, Supplementary Methods and Discussion**). Therefore, yeast surface co-expression of SsoFe2 is the preferred approach for generating magnetized yeast cell targets. Additionally, inclusion of excess yeast lacking the library’s selective nutritional marker during library screening will minimize the isolation and expansion of library cells with no specificity for the target that could be isolated from non-specific adsorption of dissociated iron oxide particles. These yeast cells can display a non-relevant protein or be non-displaying EBY100 cells. By including excess non-relevant yeast cells, it is likely that any dissociated iron oxide will complex with the non-relevant yeast cells that are in excess rather than the library yeast cells. Finally, we noticed a decrease in efficiency of yeast magnetization – both SsoFe2-mediated and non-specific – as the batch of iron oxide nanopowder aged. A potential explanation is that exposure to air causes changes in the properties of the iron oxide particles. Therefore, it helpful to test the efficiency of magnetization prior to screening libraries.

### Isolation of novel Sso7d binders to TOM22 and c-Kit using magnetic yeast expressing the target

We sought to investigate the use of magnetic yeast targets for the identification of novel binding proteins specific to the cytosolic domain of TOM22 and the extracellular domain of c-Kit. Towards that end, we screened a combinatorial library based on the Sso7d scaffold against magnetized yeast displaying the targets. Specifically, we used a combinatorial library designed by Cruz-Teran *et al* wherein the yeast cells simultaneously express the Sso7d mutants as soluble protein as well as yeast cell surface fusions^46^.

Yeast cells, co-expressing the target protein (TOM22 or c-Kit) and SsoFe2, as cell surface fusions, were magnetized by incubation with iron oxide powder as discussed earlier. The magnetized target cells were used to isolate binders from the library during four rounds of magnetic screening. Briefly, in each round of screening, magnetized cells were incubated with the library (**Figure 1**). Library cells that bound to the magnetized target cells were isolated using a magnet and expanded in selective yeast medium. The number of library and target cells was successively reduced for each round of screening. Additionally, excess EBY100 was also included to reduce the likelihood of isolating binders to native yeast surface proteins as well as to decrease the probability of isolating non-binders if iron oxide dissociated from the target cells.

The high avidity interaction between the binder and target displaying cells results in the isolation of even low affinity binders. Nevertheless, it is possible to bias selection towards high affinity binders by reducing the protein display levels on the library cells by DTT treatment, as previously described^47^. Accordingly, the Sso7d library cells were treated with DTT during the fourth round of screening to reduce the number of surface fusions on the Sso7d library cell population.

DNA isolated from the pool of binders obtained after the fourth round of screening was subjected to further random mutagenesis by error-prone PCR and used to construct a second yeast display library. This library was subjected to four rounds of screening as described earlier. Yeast displaying an irrelevant Sso7d mutant (Sso7d-hFc) was included along with EBY100 to increase the screening stringency and to minimize non-specific binding. For each target, plasmid DNA was isolated from ten individual clones recovered from the final screening round. DNA sequencing identified only two unique Sso7d mutants from the TOM22-binding population while only a single unique c-Kit binding Sso7d clone was identified (**Figure 4A, 5A**).

**Figure 4.**
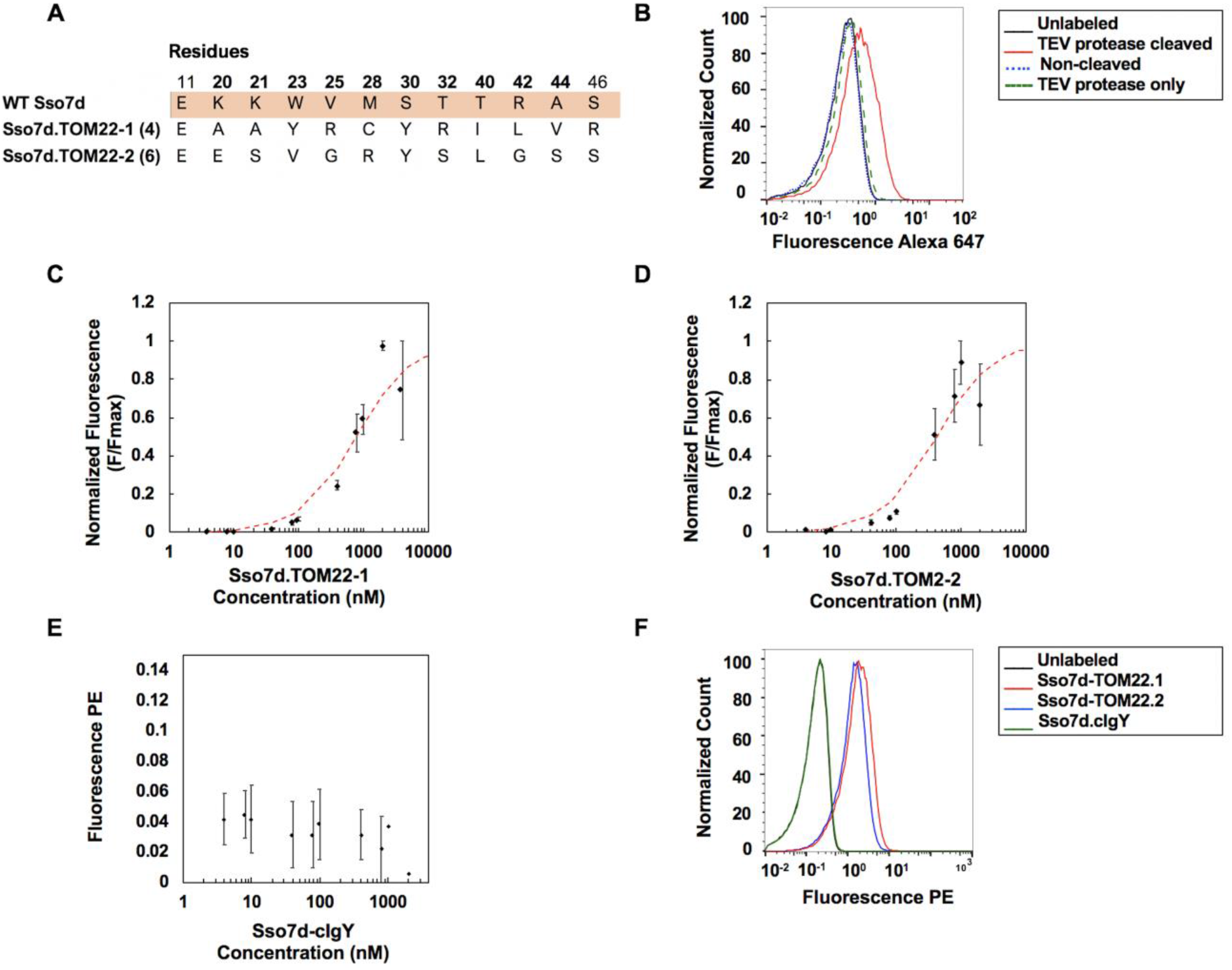
Characterization of isolated Sso7d.TOM22 mutants. (**A**) Sequences of TOM22 binders isolated from a library of Sso7d mutants. Positions mutagenized in the original library are in bold font. The number in parentheses is the number of identical DNA sequences obtained. (**B**) Soluble protein from the pool of TOM22 mutants recovered from the final round of screening was purified from yeast culture supernatant. Flow cytometry analysis was completed for unlabeled cells (black) and yeast cells expressing TOM22 upon labeling with secreted binder protein cleaved with TEV protease (red), non-treated protein (blue), and TEV protease only (green). Soluble protein (2.5 μM) and TEV protease (0.67 μM) binding was detected using an Anti-His Alexa 647 antibody. Subsequently, individual mutant analysis was completed to estimate the apparent binding K_D_ of Sso7d.TOM22-1 (**C**) and Sso7d.TOM2-2 (**D**) to yeast cells displaying TOM22. A global fit was used to estimate the K_D_ of Sso7d.TOM22-1 as 790 nM (483-1086 nM, 68% confidence interval). The K_D_ of Sso7d.TOM22-2 was estimated as 424 nM (265-626 nM, 68% confidence interval). Error bars correspond to the standard error of the mean from three independent replicates. (**E**) Background subtracted mean fluorescence values for Sso7d.cIgY binding to yeast cells displaying TOM22. Error bars correspond to the standard error of the mean from three independent replicates. Points with no error bars represent averages from two independent replicates. (**F**) Representative flow cytometry analysis of unlabeled cells (black) as well as yeast cells displaying TOM22 labeled with biotinylated Sso7d.cIgY (green), Sso7d.TOM22-1 (blue), and Sso7d.TOM22-2 (red) at a concentration of 1000 nM, followed by secondary labeling with SA-PE.

**Figure 5.**
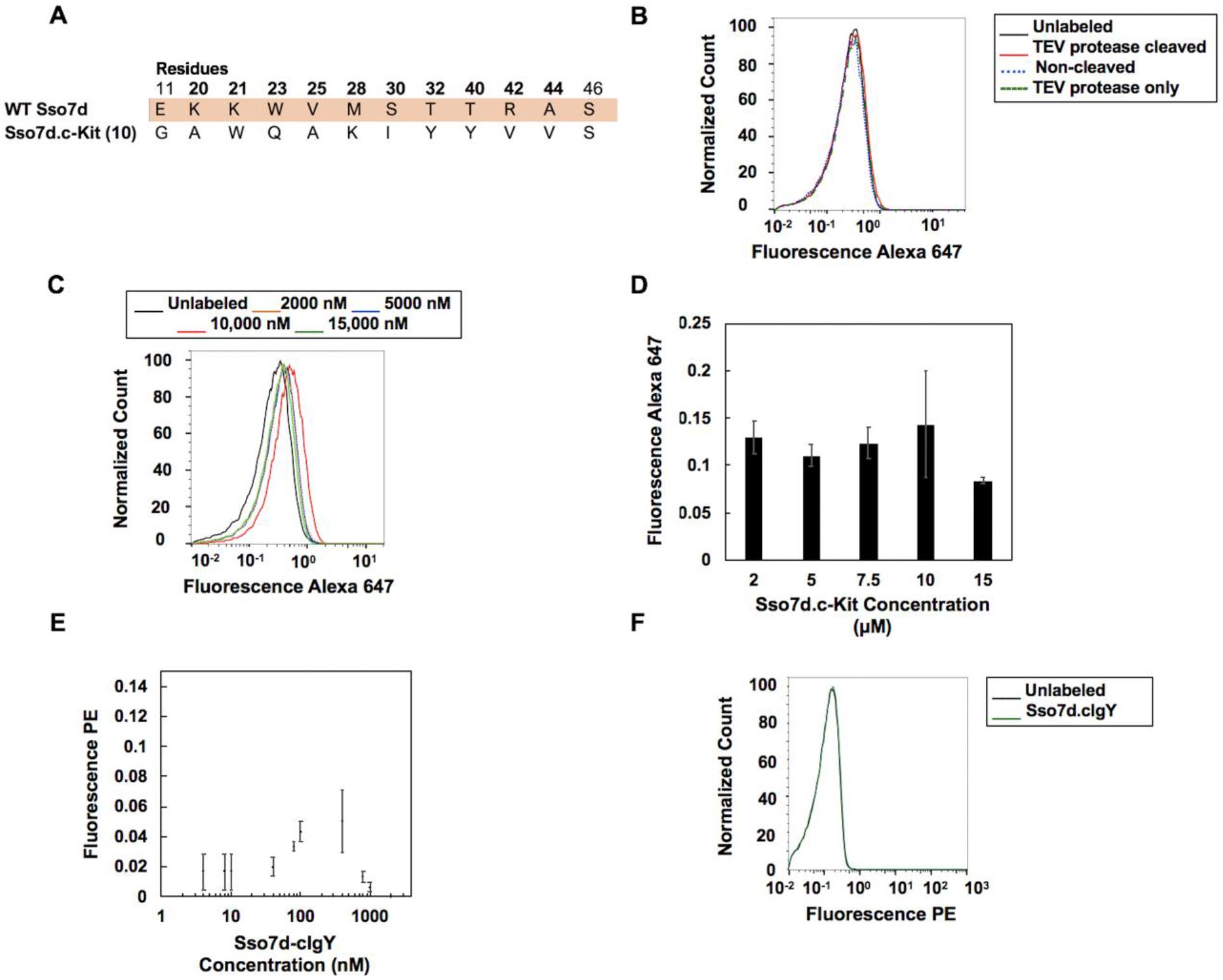
Characterization of isolated Sso7d.c-Kit mutants. (**A**) Sequences of c-Kit binders isolated from a library of Sso7d mutants. Positions mutagenized in the original library are in bold font. The number in parentheses is the number of identical DNA sequences obtained. (**B**) Soluble protein from the c-Kit mutant population recovered from the final round of screening was purified from yeast culture supernatant. Flow cytometry analysis was completed for unlabeled cells (black) and yeast cells expressing c-Kit upon labeling with secreted binder protein cleaved with TEV protease (red), non-treated protein (blue), and TEV protease only (green). Soluble protein (2.5 μM) and TEV protease (0.67 μM) binding was detected using an Anti-His Alexa 647 antibody. (**C**) Representative flow cytometry analysis of unlabeled yeast cells and c-Kit displaying yeast cells labeled with varying concentrations of Sso7d.c-Kit and an Anti-His Alexa 647 antibody. (**D**) Background subtracted mean fluorescence values for Sso7d.c-Kit binding to c-Kit displaying yeast cells at various concentrations. Error bars correspond to standard error of the mean for three independent repeats. (**E**) Background subtracted mean fluorescence values for Sso7d.cIgY binding to yeast cells displaying c-Kit. Error bars correspond to the standard error of the mean from three independent replicates. Points with no error bars represent the average of two independent replicates. (**F**) Representative flow cytometry analysis of unlabeled cells (black) as well as yeast cells displaying c-Kit labeled with biotinylated Sso7d.cIgY (green) at a concentration of 2000 nM, followed by secondary labeling with SA-PE.

### Characterization of yeast secreted Sso7d mutants

For each target, the pool of binders isolated after screening the mutagenized libraries was analyzed to confirm binding and specificity. Because a simultaneous secretion and display library was used, Sso7d binder protein could be secreted by the library yeast cells into the supernatant of the yeast display culture, without any additional sub-cloning steps. Individual clones were not characterized at this stage; rather, a mixture comprised of the final population clones was analyzed. Soluble protein, which contains a 6xHis tag, was purified from the supernatant using Ni-NTA agarose and concentrated as previously described^46^.

Notably, the soluble protein obtained from the simultaneous secretion and yeast surface display system contains a portion of a F2A peptide at its C-terminus, which can be cleaved by treatment with TEV protease. The soluble protein obtained from the TOM22 library screens bound TOM22 displaying cells when the soluble protein was treated with TEV protease to cleave the F2A peptide. No binding was observed when the F2A peptide sequence was present (**Figure 4B**). The presence of the F2A peptide sequence at the secreted protein’s C-terminus may impede the binder population from interacting with TOM22. Alternatively, the F2A peptide sequence may hinder binding of the anti-His antibody used for immunofluorescent detection. These results confirm that the pool of proteins isolated from the combinatorial library screens against yeast cells displaying TOM22 does in fact bind TOM22.

In contrast, Sso7d soluble protein derived from the c-Kit specific screens did not bind c-Kit displaying yeast cells at a protein concentration of 2.5 μM, even upon cleavage of the F2A peptide sequence by TEV protease (**Figure 5B**). These results suggest that the binding affinity of the isolated c-Kit Sso7d binder was significantly lower than that of the TOM22 Sso7d binders.

### Characterization of binding affinity and specificity of recombinantly expressed Sso7d mutants

Each identified Sso7d binder for TOM22 and c-Kit was recombinantly produced in *E. coli,* and the purified protein was used to generate yeast surface titrations to estimate the apparent K_D_ of binding. Briefly, yeast cells displaying either TOM22 or c-Kit were incubated with varying concentrations of each mutant. The fraction of cell surface fusions bound by the mutant protein was quantified by immunofluorescent detection of biotinylated protein (for TOM22 binders) or a His tag on the recombinant protein (for the c-Kit binder). Note that the recombinantly produced Sso7d.c-Kit was not biotinylated to avoid potential modification of a lysine mutation introduced in the binding interface. The data was fit to a monovalent binding isotherm, and the K_D_ was estimated as described^31^.

The apparent K_D_ of SSo7d.TOM22-1 and SSo7d.TOM22-2 was estimated as 790 nM and 424 nM respectively (**Figure 4C & 4D**). Similarly, the apparent K_D_ of Sso7d.c-Kit.1 was estimated to be greater than 2000 nM (**Figure 5C & 5D**). The exact K_D_ for Sso7d.c-Kit could not be determined due to low binding affinity; a complete yeast titration curve could not be generated. Nevertheless, weak, yet detectable levels of Sso7d.c-Kit binding to yeast displayed c-Kit was observed across concentrations ranging from 2,000 nM to 15,000 nM.

While the isolated binders have low to moderate affinity, they showed specific binding for their target. We evaluated the binding of an irrelevant Sso7d mutant, Sso7d.cIgY, to yeast displayed TOM22 and c-Kit to assess the binding specificity of the selected mutants. Sso7d.cIgY binds specifically to chicken IgY and has a reasonably similar composition of residues in its binding interface as the Sso7d TOM22 and c-Kit binding mutants^34^. Yeast surface titrations were conducted, as described earlier, for Sso7d.cIgY binding to TOM22 displaying cells (**Figures 4E & 4F**). In contrast to the TOM22-binding Sso7d mutants, Sso7d.cIgY negligibly bound to yeast displayed TOM22 over the range of concentrations tested, and the data did not fit a monovalent binding isotherm. Taken together, these results confirm that Sso7d.TOM22-1 and −2 specifically bind the yeast displayed cytosolic domain of TOM22, and this binding is not a consequence of non-specific binding of the Sso7d scaffold to TOM22. Similarly, at a concentration of 2 μM, Sso7d.c-Kit – but not Sso7d.cIgY – exhibited binding to yeast displayed c-Kit (**Figure 5C–5F**). These results show that Sso7d.c-Kit binds to yeast displayed c-Kit specifically, though with weak binding affinity (K_D_ > 2 μM).

Despite the isolation of low to moderate affinity Sso7d binders, the convergence of the Sso7d libraries to one or two sequences when selected against yeast cells displaying the target domains suggests effectiveness of this screening strategy. The low binding affinity of Sso7d.c-Kit may represent a limitation of the Sso7d scaffold for generating binders to certain targets, as was previously observed for other target proteins^48^. The binding affinity of Sso7d.c-Kit can likely be improved through additional rounds of mutagenesis and screening. However, such affinity maturation was not pursued since the focus of our study was investigating the feasibility of screening yeast display libraries using magnetized yeast cell targets. Instead, we chose to extend our results by screening a combinatorial library based on a different scaffold protein.

### Isolation of nanobodies specific to TOM22 & c-Kit

We investigated the screening of a combinatorial library based on a camelid single-domain antibody fragment (or nanobody) scaffold, as described in McMahon *et al*^35^, against magnetized yeast displaying a target of interest. In contrast to single chain antibody fragments, nanobodies have a single 15 kDA V_HH_ domain. The nanobody library was screened to obtain binders to TOM22 and c-Kit using magnetized yeast cell targets, as described earlier. The selection stringency was increased in successive rounds of screening. Note, however, that the nanobody library cannot be treated with DTT to reduce the surface fusion copy number. This is because the nanobody mutants are covalently tethered to the cell wall by a synthetic amino acid chain designed to mimic low complexity yeast cell wall proteins, as opposed to being tethered by disulfide bonds to the cell wall via the interaction between Aga2 and Aga1, like the Sso7d library.

For each target, plasmid DNA was isolated from ten individual clones recovered during the final screening rounds. DNA sequencing identified 8 unique mutants from the TOM22 and c-Kit nanobody binding populations (**Figure 6A, B**). For each target, two unique mutants were recombinantly produced in *E.coli,* and the purified protein was biotinylated. The mutants chosen for characterization appeared twice in the sequenced populations. For each mutant, the apparent K_D_ of binding to its target was measured using yeast surface titrations in a similar manner as described for the Sso7d mutants. The apparent K_D_ of NB.TOM22-1 and NB.TOM22-2 (**Figure 6C**) was estimated as 271 and 2009 nM, respectively, while the apparent K_D_ of NB.c-Kit-1 and NB.c-Kit-2 (**Figure 6D**) was estimated as 131 and 204 nM, respectively. Except for NB.TOM22-2, the binding affinities of the isolated nanobody mutants were higher than the corresponding Sso7d mutants, underscoring differences between these scaffolds. Nevertheless, isolation of NB.TOM22-2 confirms that even binders with low affinity can be isolated when magnetized target cells are used.

**Figure 6.**
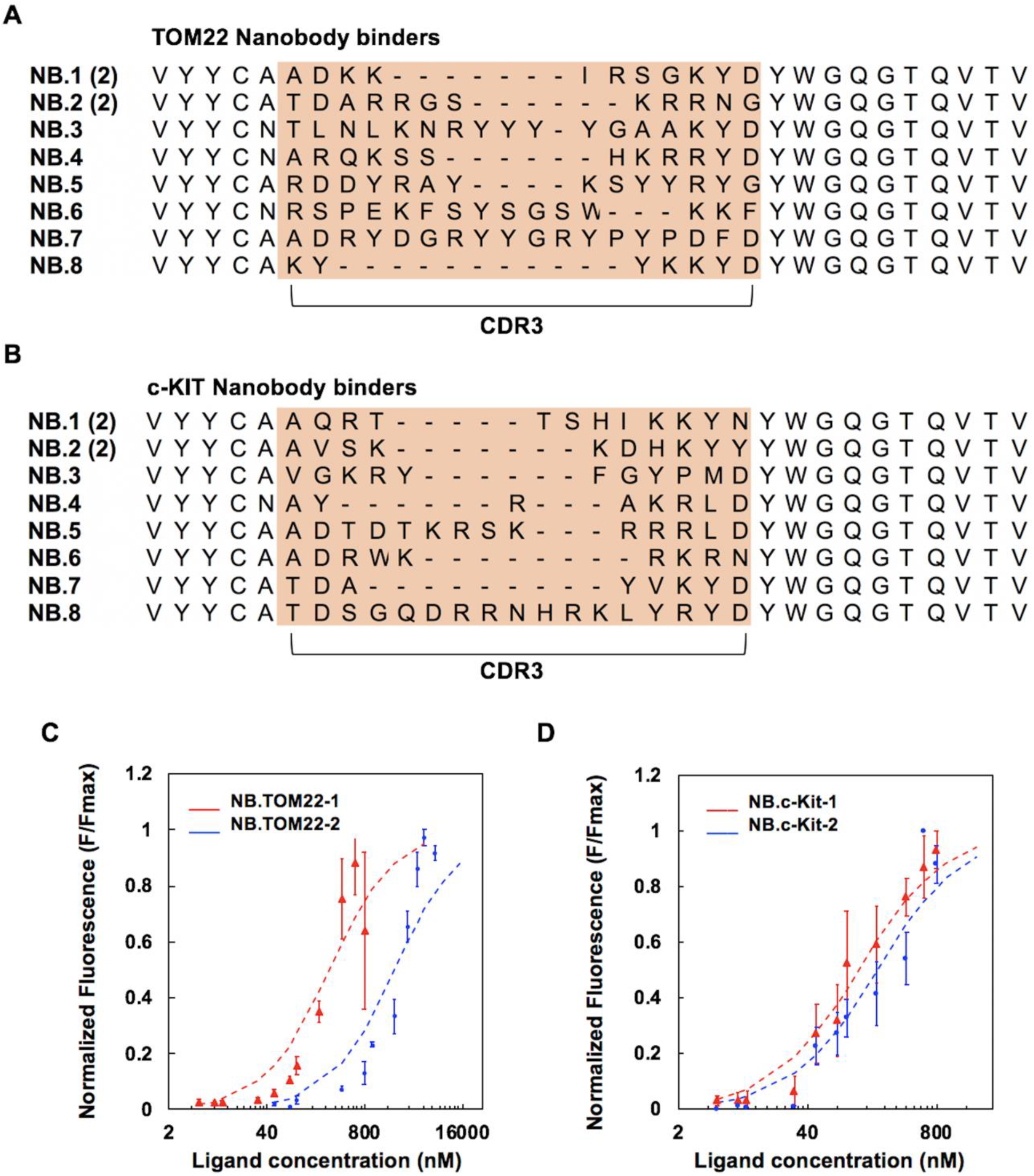
Analysis of isolated nanobody binders. Sequences of (**A**) TOM22 binders and (**B**) c-Kit binders isolated from a nanobody combinatorial library. CDR3 positions are highlighted in light brown. The number in parentheses is the number of identical DNA sequences obtained. (**C**) Yeast surface titrations to estimate the apparent K_D_ of NB.TOM22-1 (red) and NB.TOM22-2 (blue) for TOM22. A global fit was used to estimate the K_D_ for NB.TOM22-1 as 271 nM (220-338 nM, 68% confidence interval). The K_D_ for NB.TOM22-2 was estimated as 2009 nM (1504-2690 nM, 68% confidence interval). (**D**) Yeast surface titrations to estimate the apparent K_D_ of NB.c-Kit-1 (red) and NB.c-Kit-2 (blue) for c-Kit. The K_D_ for NB.c-Kit-1 and NB.c-Kit-2 was estimated as 131 nM (116-149 nM, 68% confidence interval) and 204 nM (166-250 nM, 68% confidence interval), respectively, using a global fit. Error bars correspond to the standard error of the mean from three or four independent replicates.

To further test the functionality of the isolated c-Kit binders, c-Kit-specific nanobodies were immobilized on magnetic beads and evaluated for their ability to deplete yeast cells expressing c-Kit from a heterogeneous population. However, the c-Kit displaying cells were only depleted around 10% after incubation with magnetic beads functionalized with NB.c-Kit-1 or NB.c-Kit-2 (**Figures S3, S4; Supplementary Methods and Discussion**). It is likely that a higher affinity binder is needed to achieve significant depletion of c-Kit displaying cells. Therefore, we did not pursue further characterization of the c-Kit binding nanobodies in the context of isolating mammalian cells that naturally express c-Kit from a heterogenous mixture.

### TOM22 binders can enrich mitochondria from cell lysates via recognition of natively expressed TOM22

To characterize the functionality of the TOM22 binders, we assessed the isolated TOM22 Sso7d and nanobody binders for their ability to recover mitochondria from a cell lysate. Each TOM22 binder protein was recombinantly produced, biotinylated, and immobilized onto streptavidin-functionalized 0.15 μm beads. Subsequently, lysates from HEK293-T cells were incubated with the binder-coated beads, and the beads were recovered. Immunoblotting was used to quantify the recovery of mitochondria pulled down by the binder-functionalized beads. Mitochondrial recovery was assessed by immunoblotting with an anti-TOM22 antibody (**Figure 7A**). Pulldown of a non-specific organelle, the endoplasmic reticulum, was assessed using an anti-calnexin antibody (**Figure 7B**); this serves as a proxy for the interaction of the binders with non-target species. Commercially available micrometer-sized magnetic beads functionalized with an anti-TOM22 antibody are commonly used for the separation of mitochondria from cell lysates^49–52^. We carried out isolation of mitochondria from HEK293-T lysates using the commercially available antibody coated magnetic beads in parallel. Results from these beads served as a benchmark to evaluate the efficiency of the TOM22 binders identified in this study.

**Figure 7.**
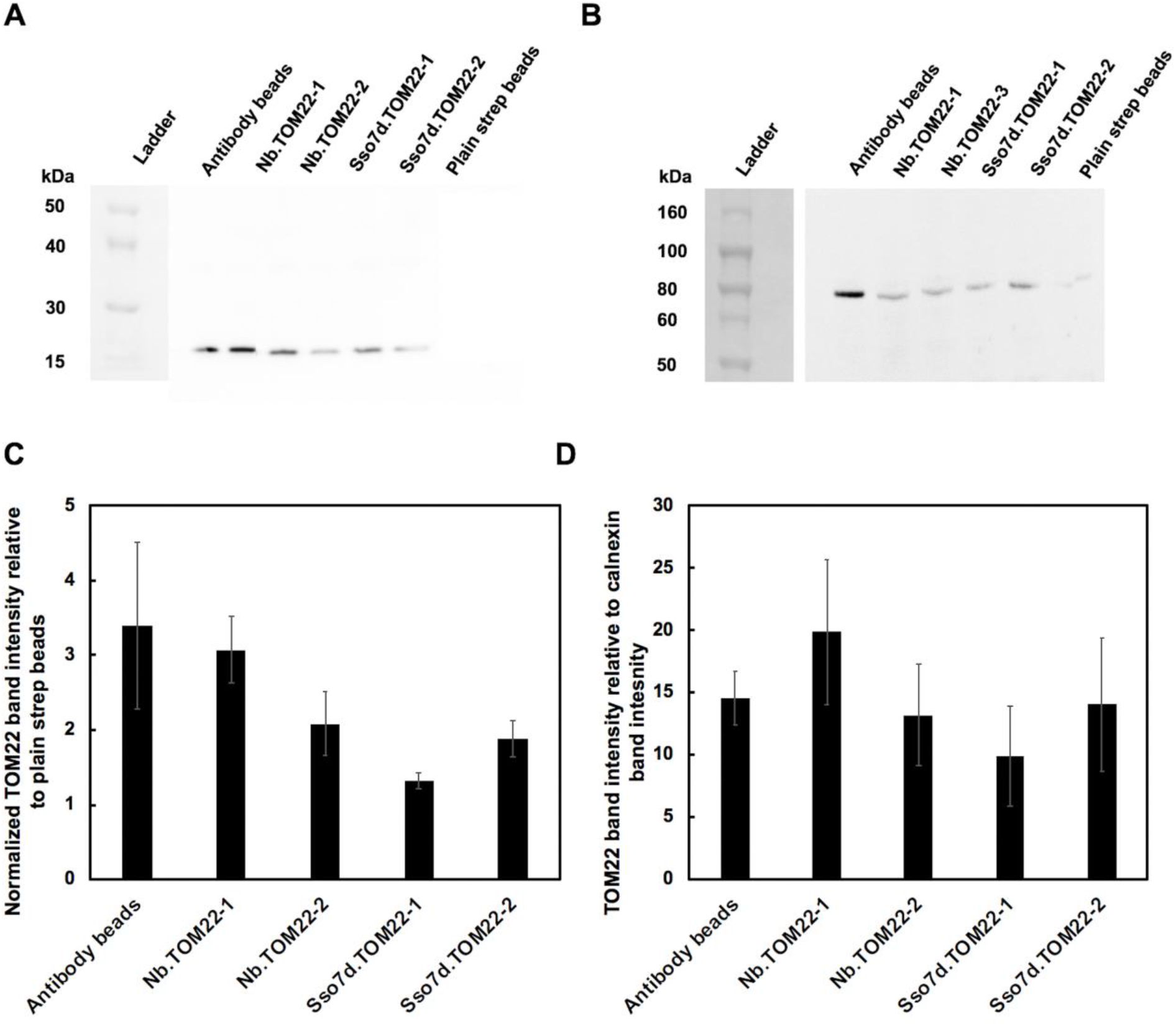
Mitochondria enrichment for the identified TOM22 nanobody and Sso7d binders. The binder proteins were biotinylated and functionalized onto streptavidin beads (0.15 μm). The mitochondrial recovery of the TOM22 nanobody and Sso7d binders was compared to the recovery observed using non-functionalized streptavidin beads as well as commercially available micro-sized magnetic beads functionalized with an anti-TOM22 antibody. (**A**) Mitochondrial recovery was quantified using anti-TOM22-HRP immunoblotting. (**B**) Endoplasmic reticulum recovery was quantified using anti-Calnexin-HRP immunoblotting. An image of the ladder was obtained in a separate image than the blot. Blot images were inverted for display. The ladder and experimental condition sections of the blot were spliced together after alignment. No other manipulations of the image occurred. These blots are representative of three replicates. Mitochondria and endoplasmic reticulum recovery was quantified by band intensity analysis. (**C**) The mitochondrial recovery by beads functionalized with a nanobody, Sso7d, or antibody specific for TOM22 was compared to the mitochondrial recovery of non-functionalized streptavidin beads to evaluate the enrichment of each binding protein against background binding. (**D**) The specificity of the binder proteins was evaluated by comparing the mitochondrial recovery to the endoplasmic reticulum recovery for each bead type considered. Error bars represent the standard error of the mean for at least three independent replicates.

We observed that the mitochondrial recovery, as assessed by immunodetection of TOM22, was higher when each mutant was immobilized onto the streptavidin beads in comparison to the recovery using plain (unmodified) beads (**Figure 7C**); NB.TOM22-1 exhibited the greatest recovery among the identified ligands. A direct comparison of mitochondrial recovery between the beads coated with Sso7d or nanobody binders and the commercial product is not possible since the surface density of the anti-TOM22 antibody is unknown. However, we used the ratio of the TOM22 band intensity to that of the calnexin band intensity as a quantitative metric for specificity, to compare the commercial beads with the binder-coated streptavidin beads. Based on this metric, our mutants have specificity for mitochondria over the endoplasmic reticulum comparable to that of the commercial antibody beads (**Figure 7D**). The Sso7d and nanobody mutants were able to recover mitochondria from the heterogenous cell lysate with similar efficiency as the antibody coated beads. These results suggest the Sso7d and nanobody binders specific to TOM22, isolated from selections against a yeast displayed target, are functional in the context of the native mitochondrial protein.

Increases in yield and specificity may be possible if the pull-down protocol is further optimized. However, we did not optimize the binding and wash conditions. Furthermore, our pull-down protocol with the streptavidin coated beads used centrifugation to isolate the beads as opposed to magnetic separation, which was used to recover the commercial beads. It is also noteworthy that NB.TOM22-2 showed similar or greater mitochondria recovery in comparison to the Sso7d mutants, despite its weak affinity ~2000 nM. A potential explanation is that protein immobilization may differentially affect the binding activity of the various mutants. Alternatively, these differences may be attributed to increased non-specific binding of proteins present in cell lysates to the Sso7d mutants. While the specificity of the ligands for mitochondrial capture over endoplasmic reticulum was explored, there are a wide variety of other proteins and organelles found in the cell lysate that were not considered when evaluating specificity that could be outcompeting the mitochondria for binding to the functionalized beads. Therefore, inclusion of target-depleted cell homogenates during later rounds of library screening may be beneficial for isolation of binders with greater specificity for TOM22. Nevertheless, our results confirm that our isolated Sso7d and nanobody mutants bind natively expressed TOM22 on the surface of a mitochondria.

In conclusion, we have shown that the isolation of binding proteins from yeast display libraries using yeast displayed targets is feasible. A critical aspect of our strategy is affinity-based magnetization of target-displaying cells, mediated by interactions between iron oxide nanoparticles and a co-expressed iron oxide binding protein; this facilitates facile, magnetic separation of library cells that bind the target. Binding proteins discovered when selecting against yeast displayed targets were functional in the context of a natively expressed membrane protein, as seen with the TOM22 binders depleting cell lysates of mitochondria. We expect that this strategy will enable efficient isolation of binders to membrane protein targets.

## Materials and Methods

### Plasmids and yeast culture

The *Sacchromyces cerevisiae* strain EBY100 was used in conjunction with the pCTCON vector containing a *TRP* selectable marker, or a pCT302 vector variant with a *LEU* selectable marker^53^. Plasmid DNA was transformed into chemically competent EBY100 using the Frozen-EZ yeast transformation Kit II (Zymo Research).

For cells with the pCTCON vector, Trp-deficient SDCAA and SGCAA medium was used for culturing cells and for inducing cell surface protein expression, respectively, as previously described^31^. Similarly, Leu-deficient SDSCAA (-Leu) and SGSCAA (-Leu) media was used for cells with the pCT302-based vector. Leu-deficient media has similar composition to SDCAA and SGCAA media except it contains synthetic dropout mix (1.62 g/L) lacking leucine instead of casamino acids. Yeast cells were cultured in SDCAA or SDSCAA medium, as appropriate, at 30°C with shaking at 250 RPM. Protein expression was induced by transferring the yeast cells into SGCAA or SGSCAA medium at an OD_600_ of 1 and cultured overnight at 20 °C with shaking at 250 rpm. EBY100 without plasmid was grown in YPD medium (10.0 g/L yeast extract, 20.0 g/L peptone, and 20.0 g/L dextrose) at 30 °C with shaking at 250 rpm.

The nanobody library was grown in TRP deficient NB.SDCAA and NB.SGCAA medium (3.8 g −TRP drop-out media supplement (US Biological), 6.7 g Yeast nitrogen base (HiMedia), 20 g dextrose for growth or galactose for protein induction, pH 6). The nanobody library cells were cultured in NB.SDCAA medium at 30 °C with shaking at 250 RPM. Protein expression was induced by transferring the yeast cells into NB.SGCAA medium at an OD_600_ of 1 and cultured overnight at 20 °C with shaking at 250 rpm.

### Construction of co-expression plasmids

pCTCON-SsoFe2-T2A-TOM22 was constructed by amplifying gene block 1 with Pf1 and Pr1 and introduced to the pCTCON vector between the EcoRI and XhoI sites. The DNA corresponding to the Fc portion of human IgG (residues 100-330 of immunoglobulin heavy constant gamma 1) was amplified from gene block 2 using primers Pf2 and Pr2 and inserted into pCTCON-T2A-SsoFe2-TOM22 between the NheI and BamHI sites creating pCTCON-T2A-SsoFe2-hFc.

To construct pCT302-SSoFe2-T2A-TOM22, the pCT302 vector was cut between NheI and XhoI. Gene block 1 was amplified by PCR using primers Pf1 and Pr1 and introduced into pCT302 using homologous recombination. Briefly, 200 ng of cut plasmid and 250 ng of insert were transformed into EBY100 cells prepared with the Frozen-EZ yeast transformation kit II. The plasmid and insert were concentrated using ethanol precipitation such that the volume of DNA used in the transformation step was less than 5 μL in total. The extracellular domain of c-Kit (residues 2-83) was amplified from gene block 3 using primers Pf3 and Pr3. pCT302-SsoFe2-T2A-c-Kit was constructed by inserting the amplified c-Kit DNA between the NheI and BamHI sites in pCT302-SsoFe2-T2A-TOM22. pcT302-SsoFe2-T2A-hFc was constructed in a similar manner by amplifying the hFc DNA from gene block 2 with primers Pf2 and Pr2 and inserting between the NheI and BamHI sites of pCT302-SsoFe2-T2A-TOM22.

To generate the vector where simultaneous display is mediated by the bidirectional *GAL1/GAL10* promoter, gene block 4 was amplified using primers Pf4 and Pr4 and introduced to the pCTCON vector between the EcoRI and XhoI sites to eliminate an AgeI site from the prepro sequence used in previous constructs to generate pCTCON-modified-prepro. Gene block 5 was amplified using primers Pf5 and Pr5 and inserted into the pCTCON-modified-prepro plasmid between the KpnI and AgeI sites creating pCTCON-SsoFe2-Bi-TOM22. DNA corresponding to hFc was amplified using primers Pf2 and Pr2 and inserted into pCTCON-SsoFe2-Bi via the NheI and BamHI sites to generate pCTCON-SsoFe2-Bi-hFc.

Sso7d-hFc was inserted between the NheI and BamHI sites of pCTCON and pCT302 to create pCTCON-Sso7d-hFc and pCT302-Sso7d-hFc^44^. The Sso7d-hFc DNA was amplified using primers Pf6 and Pr6 from previously constructed plasmids^44^. The TOM22 mitochondrial domain was cloned into pCT302 by amplifying gene block 1 using primers Pf7 and Pr7 and inserting the amplified, digested DNA between the NheI and the BamHI sites to create pCT302-TOM22. The extracellular domain of c-Kit was similarly cloned into pCT302 using gene block 3 and primers Pf3 and Pr3 to generate pCT302-c-Kit.

All gene fragments were purchased from Integrated DNA technologies. Primer oligonucleotides were purchased from IDT or Eton Biosciences. Gene fragment and primer sequences can be found in **Supplementary Tables S1 and S2**. All PCR reactions were performed in 50 μL reactions with high-fidelity Phusion polymerase (Thermo Fisher Scientific) according to the manufacturer’s protocol. All restriction enzymes were purchased from New England Biolabs. Restriction digestions were performed in 50 μL with a 5 times excess of each restriction enzyme for 4 hours at 37°C. Digested plasmid backbones were treated with Antarctic Phosphatase purchased from New England Biolabs for 1 hour at 37°C. Digested products and PCR amplicons were purified using the 9K Series Gel and PCR extraction kit from Biobasic. Ligations of the digested plasmid backbones and PCR products occurred overnight at 16°C using T4 DNA ligase (Promega) prior to transformation into chemically competent Novablue *E. coli* cells. The Novablue cells were made chemically competent using the Mix & Go! E. coli transformation kit and buffer (Zymo Research). Overnight *E. coli* cultures were harvested for their plasmid using the GeneJET plasmid miniprep kit (Thermo Fisher Scientific).

### Expression analysis of dual display constructs

The expression level of the co-expressed proteins for both dual display constructs was estimated using flow cytometry analysis using pCTCON-T2A-SsoFe2-hFc and pCTCON-SsoFe2-Bi-hFc. Briefly, 5×10^6^ cells were labeled with a 1:100 dilution of chicken anti-c-myc antibody or mouse anti-V5 antibody (Thermo Fisher Scientific) for 15 minutes at room temperature. Subsequently, cells were washed and secondary labeling was carried out using a 1:250 dilution of goat-anti-chicken 633 or donkey anti-mouse 633 (Immunoreagents, Raleigh, NC) for 10 minutes on ice. All labeling was conducted in 50 μL of 0.1% PBS BSA (PBSA). Cells were analyzed using a Miltenyi Biotec MACsQuant VYB cytometer.

### Magnetic pull downs of a known binder-target pair

5×10^7^ pCTCON-T2A-SsoFe2-hFc yeast cells and 5×10^7^ cells pCT302-T2A-SsoFe2-hFc yeast cells were spun down in separate tubes and resuspended in 2 mL of 1% BSA TN buffer (50 mM Tris-HCl, 300 mM NaCl pH 7.4; TNBSA). To magnetize the cells, 100 μL of a 4 mg/mL iron oxide (II, III) (Thermo Fisher Scientific) solution in water was added to the cells and incubated for 30 minutes. The non-magnetized cells were removed using a magnet. OD_600_ of the solution prior to the addition of iron oxide (OD_i_) and after the removal of yeast bound to iron oxide (OD_f_) was measured using 100 μL of the sample. The number of cells bound to the iron oxide was calculated as (OD_i_-OD_f_) and used for normalization between replicates, as described later. The cells complexed with the iron oxide were washed three times with 1% TNBSA and blocked with 2 mL of 1% TNBSA for 1 hour. After blocking, the cells complexed with the iron oxide were washed two times with 0.1% TNBSA.

Varying amounts of pCT302-TOM22 yeast cells and pCTCON-Sso7d-hFc yeast cells were incubated with the magnetized target cells (10^7^:10^6^, 10^8^:10^6^, 10^9^:10^6^, 10^9^:10^5^, and 10^9^:10^4^ pcT302-TOM22 cells: pCTCON-Sso7d-hFc cells) for two hours at room temperature in 2 mL of 0.1% TNBSA with rotation. Two separate cell aliquots were generated for each ratio. One aliquot was incubated with magnetic target cells that contained the pCTCON-T2A-SsoFE2-hFc plasmid (*TRP* selectable marker) while the other aliquot was incubated with magnet target cells that contained the pCT302-T2A-SsoFe2-hFc plasmid (*LEU* selectable marker). At the end of the incubation period, the supernatants containing the unbound cells were removed after placing the mixtures on a magnet, and the cells complexed with the iron oxide were washed three times with 0.1% TNBSA. The iron oxide and any bound cells were resuspended in 1 mL of 0.1% TNBSA to generate a final sample mixture.

100 μL of 1/10, 1/100, and 1/1000 dilutions of the final sample mixture were plated. Samples that used magnetic target cells containing the pCTCON-T2A-SsoFe2-hFc plasmid were plated on SD (-Leu) plates to quantify non-specific, TOM22 yeast binding to the target hFc yeast cells. Samples containing magnetic target cells based on the pCT302-T2A-SSoFe2-hFc plasmid were plated on SD (-Trp) plates to quantify specific, Sso7d-hFc yeast binding to the target hFc yeast cells (**Figure S1**). Colony counting was used to determine the number of Sso7d-hFc and TOM22 yeast recovered by the magnetic target cells. Ultimately, these values were used to quantify enrichment and recovery of the Sso7d-hFc binder populations. There were slight differences in the number of cells magnetized for each replicate. The number of captured cells was scaled by the number of magnetized cells for each replicate.

Additionally, 1×10^6^ pCTCON-Sso7d-hFc and pCT302-TOM22 cells were spun down and resuspended in 1 mL of 0.1% TNBSA. 100 μL of a 1/1000 dilution of these cells were plated on -Trp and -Leu plates, respectively, and counted to quantify the number of these cells initially incubated with the magnetized target cells. Enrichment is defined as the percentage of Sso7d-hFc cells in the final population compared to the percentage of Sso7d-hFc cells in the initial population. Recovery is defined as the percentage of Sso7d-hFc cells from the initial population that were isolated in the population isolated by the magnetic target cells.

### Sso7d library screening using targets displayed on magnetic yeast

Multiple rounds of magnetic sorting were carried out to isolate binders to TOM22 and c-Kit. A simultaneous secretion and display library of Sso7d mutants (-Trp) was used ^46^. The library was screened against pCT302-T2A-SsoFe2-TOM22 (-Leu) or pCT302-T2A-SsoFe2-cKit (-Leu).

In the first round, 5×10^8^ cells expressing the target and SsoFe2 were resuspended in 9 mL of 1% TNBSA. The cells were magnetized by adding 1 mL of an iron oxide solution (4 mg/mL) followed by a 30 minute incubation at room temp. Based on the OD_600_ of the solution prior to the addition of the iron oxide and after the removal of yeast bound to the iron oxide, 2.5×10^8^ magnetized cells were aliquoted for use in the magnetic selections. The magnetized cells were washed 3X with 1% PBSA buffer and blocked with 1% PBSA for 1 hour at room temp in a 10 mL volume. After, the cells were washed 3X with 0.1% PBSA prior to the addition of 2×10^9^ library cells and 2×10^9^ EBY100 cells in 10 mL of 0.1% PBSA. The selection took place for 2 hours at room temperature. Library cells bound to the magnetized target cells were separated from the unbound cells using a magnet. The iron oxide was washed 3 times with 0.1% PBSA prior to expansion in 20 mL of SDCAA (-TRP) media.

A similar process was carried out in the subsequent magnetic selection rounds. However, there was a reduction in number of magnetized target cells and library cells for these rounds. In the second round, 2.5×10^7^ magnetic target cells were incubated with 1×10^7^ library cells and 1×10^9^ EBY100 cells. The target-bound library cells were expanded in 5 mL of SDCAA (-Trp) media. In the third round, 2.5×10^6^ magnetic target cells were incubated with 1×10^5^ library cells and 1×10^9^ EBY100 cells. In these rounds, magnetization of the target cells took place in a total volume of 2 mL by incubating 5×10^7^ or 5×10^6^ cells in 1% PBSA, respectively for rounds 2 and 3. For round 2, 100 μL of the 4 mg/mL iron oxide solution was added while 10 μL of the iron oxide solution was incubated for round 3. After magnetization, the cells were appropriately aliquoted to carry forward the correct number of magnetized target cells.

In the fourth round, the library cells were treated with DTT to reduce display levels by ~80%, similar to Stern *et al*^47^. The DTT concentration resulting in a ~80% reduction of c-myc expression was empirically determined for each library using the following procedure. 5×10^6^ cells from the third round of the library screening were pelleted and washed twice with 10 mM Tris pH 7.5 buffer (1.24 g/L Tris-HCl, 0.26 g/L Tris base). Solutions of varying DTT concentration (1, 5, 7.5 10, 15, or 20 mM) were created using the 10 mM Tris pH 7.5 buffer. The yeast cells were resuspended in 20 μL of the different DTT concentration solutions at 30°C for 20 minutes with shaking. After, the cells were pelleted and washed twice with 0.1% PBSA. To quantify ligand expression, the treated yeast cells were labeled with 50 μL of a 1:100 dilution of chicken-anti-c-myc antibody for 15 minutes at room temperature, then washed once with 0.1% PBSA. Secondary labeling was performed using 50 μL of a 1:250 dilution of goat-anti-chicken 633 on ice. The mean fluorescence of 50,000 events from each sample was compared to an untreated control to determine the reduction in the display level. It was determined that 5 mM and 7.5 mM of DTT was required to reduce the display levels of the putative TOM22 and c-Kit binders by ~80%, respectively. During the fourth round of selection, 1×10^5^ DTT treated library cells and 1×10^9^ EBY100 were incubated with 1×10^6^ magnetized target cells. The target cells were magnetized as previously described for round 3 and appropriately aliquoted.

### Construction of a second Sso7d yeast display library by random mutagenesis using error-prone PCR

DNA associated with the mutants recovered from the fourth round of screening was isolated using a Zymoprep yeast plasmid miniprep II kit (Zymo Research). For both the TOM22 and c-Kit screens, the Sso7d sequences from the isolated DNA were amplified by error prone PCR using nucleotide analogs with primers Pf6 and Pr6, as detailed by Gera *et al*^44^. The amplified error-prone PCR DNA was combined and amplified further using primers Pf8 and Pr8 in six identical 50 μL reactions. The PCR products were combined and purified using phenol: chloroform extraction. The DNA was concentrated using ethanol precipitation. Briefly, 1/10 volume of potassium acetate was added to the extracted DNA along with 10 μL of linear acrylamide followed by 2 volumes of ice-cold ethanol. The mixture was incubated at −20 °C overnight and centrifuged at 15,000 g for 10 minutes followed by removal of the supernatant. The pellet was washed once with 70% ethanol followed by a 100% ethanol wash and allowed to dry prior to resuspension in water. In parallel, 40 μg of pCT-NT-F2A-Sso7dhFc from Cruz-Teran *et al.* was digested with NheI and BamHI restriction enzymes and concentrated by phenol:chloroform extraction and ethanol precipitation^46^. A yeast display library was generated by transforming the PCR product and the digested vector into electrocompetent yeast using the lithium acetate method previously described^54^. Two electroporation reactions were performed using a Bio-Rad Gene Pulser system (Bio-Rad) where 12 μg of PCR product and 4 μg of digested vector was added to 400 μL of electrocompetent *S. cervisiae strain* EBY100. The electroporation was carried out at 2500 V, 25 μF, and 250 Ω. As a control, an identical electroporation reaction containing only the digested vector was also performed. The library diversity was quantified by plating serial dilutions of the transformation reaction onto SDCAA plates and estimated as 6×10^7^ for the TOM22-binder library and 5×10^7^ for the c-Kit-binder library.

### Additional screening of mutagenized Sso7d library

Four additional rounds of magnetic selection were carried out using the libraries constructed by error prone PCR. Yeast cells dually displaying SsoFe2 and TOM22 or SsoFe2 and c-Kit were used as targets. For rounds 1-3, 1×10^9^ EBY100 and 1×10^9^ pCT302-Sso7d-hFc cells were incubated along with the library cells. All selections took place in 0.1% PBSA. For round 1, 6×10^8^ and 5×10^8^ library cells were incubated for the TOM22 and c-Kit screens, respectively, in a volume of 10 mL. 3×10^8^ cells with pCT302-TOM22-T2A-SsoFe2 or pCT302-c-Kit-T2A-SsoFe2 were magnetized. All magnetization and incubation steps were carried out as previously described. In round two, 1×10^7^ library cells were incubated with 2.5×10^7^ magnetized target cells while in round three 1×10^6^ library cells were incubated with 2.5×10^6^ magnetized target cells in a volume of 2 mL total. In the fourth round, the library cells were treated with 7.5 mM DTT as previously described to reduce the surface display level. 1×10^5^ DTT treated library cells were incubated with 1×10^6^ magnetized target cells in a volume of 2 mL. A Zymoprep yeast plasmid miniprep II kit (Zymo Research) was used to extract the plasmid DNA of the clones recovered in the final populations. After, the DNA was transformed into electrocompetent Novablue cells and plated. 10 colonies were sequenced per target.

### Nanobody library screening against targets displayed on magnetic yeast

The nanobody yeast display library used was gifted by the Kruse lab at Harvard University^35^. This library was screened against pCT302-T2A-SsoFe2-TOM22(-Leu) or pCT302-T2A-SsoFe2-c-Kit (-Leu) yeast cells for multiple rounds.

During the first screening round, a negative selection against pCT302-T2A-SsoFe2-hFC cells was performed prior to carrying out a positive selection against cells expressing either TOM22 or c-Kit. 2.5×10^8^ magnetized cells were used in both the negative and positive selection steps. Magnetization took place in 11.5 mL of 1% PBSA for 5×10^8^ cells using 2.6 mL of a 4 mg/mL iron oxide solution. The magnetized cells were appropriately aliquoted to carry forward the correct number of magnetized cells. After, the magnetized cells were blocked in 10 mL of 1% PBSA for one hour prior to their use and subsequently washed. Next, 5×10^9^ nanobody library cells in 10 mL of 0.1% PBSA were incubated with the hFc displaying magnetized cells for one hour at room temp. After, the selection tube was placed on a magnet. Any cells that did not bind to the hFc cells were removed and incubated with the target displaying magnetized cells for 1 hour at room temperature. Finally, the iron oxide was washed four times with 0.1% PBSA prior to expansion in 20 mL of NB.SDCAA (-TRP) media.

A similar process was performed for subsequent rounds of magnetic selection with a reduction in the number of magnetized target cells and library cells. Negative and positive selection as previously described were carried out for each round. In the second round, 2.5×10^7^ magnetic cells were incubated with 5×10^7^ library cells and 1×10^9^ EBY100 cells in 2 mL of 1% PBSA + 0.05% Tween-20. The target-bound library cells were expanded in 5 mL of NB.SDCAA (-TRP media). In the third round, 1×10^6^ magnetic target cells were incubated with 5×10^5^ library cells, 1×10^9^ EBY100 cells, and 1×10^9^ pCT302-Sso7d-hFc cells in 2 mL of 1% PBS BSA + 0.05% Tween-20. No negative selection took place for the third round. In rounds two and three, the magnetization of the hFc displaying cells as well as the target cells took place in 2 mL of 1% PBSA using 200 μL of a 4 mg/mL iron oxide solution and 5×10^7^ cells. The magnetized cells were appropriately aliquoted to ensure the correct number of magnetized cells were carried forward. For both rounds, the iron oxide was washed five times with 1% PBS BSA + 0.05% tween with 10 seconds of vortexing in between washes. A Zymoprep yeast plasmid miniprep II kit was used to recover the plasmid DNA of the clones isolated in the final populations. The DNA was transformed into electrocompetent Novablue cells and plated. 10 colonies were sequenced per target.

### Specificity analysis of secreted Sso7d binder populations

For the Sso7d screens, soluble protein representing the entire pool of binders isolated from the final selection was obtained by inducing the isolated cell populations for 72 hours in SGCAA medium 30°C. The secreted protein was purified from the supernatant using Ni-NTA agarose resin (Qiagen) as previously described^46^. The purified protein was concentrated using a Vivaspin 6, 3 kDa MWCO concentrator (Sartorius), and the protein concentration was estimated using a BCA assay.

Binding of the secreted pool to its respective target was analyzed using flow cytometry. The secreted TOM22 and c-Kit binders were incubated with pCT302-TOM22 and pCT302-c-Kit yeast cells, respectively. The secreted protein was treated overnight at 4°C with TEV protease to cleave the c-myc tag and F2A sequence from the protein. 1 μg of TEV protease was incubated per 25 μg of Sso7d protein. 2.5 μM of TEV protease-treated ligand was incubated with 5×10^6^ target cells for 1 hour at room temperature. The cells were washed 1X with 0.1% PBSA prior to the incubation. The TEV protease and the secreted protein both contain 6×His tags. Therefore, target cells were also incubated with TEV protease alone to assess non-specific binding of the protease. The TOM22 and c-Kit cells were labeled with 0.046 μM of TEV protease in 50 μL of 0.1% PBSA, respectively. The target cells were also labeled with 2.5 μM of untreated, secreted binder protein. After these incubations, the cells were washed once with 0.1% PBSA and incubated with 50 μL of a 1:100 dilution of anti-penta-His Alexa 647 antibody (Qiagen) for 10 minutes on ice. A secondary only control was also performed. After a final wash, the cells were analyzed using flow cytometry. The estimated molecular weight of the Sso7d mutant population is 13 kDa.

The TEV protease was expressed in Rosetta™ *E. coli* cells using the plasmid pRK793 (Addgene, Plasmid # 8827)^55^. Cells expressing the TEV protease were grown using the following protocol. A 5 mL overnight culture was grown in LB media and ampicillin and added to 1 liter of LB media that also contained 1X M9 minimal media, 1X 5052, 1 mM MgSO_4_, and 1X ampicillin. 50X M9 minimal media contains 33.9 g/L Na_2_HPO_4_, 15 g/L KH_2_PO_4_, 5 g/L NH_4_Cl, 2.5 g/L NaCl while 50X 5052 is 25% glycerol, 2.5% glucose, 10% alpha-D-lactose. The cells were grown at 37°C until an OD_600_ of 1 was reached. After, the cells were grown at 18 °C and harvested 24 hours later. The protein was purified using a BioLogic LP FPLC system (Bio-Rad) by metal affinity chromatography. The cells were collected and lysed via sonication in 35 mL of Buffer A-IMAC (50 mM Tris, 300 mM NaCl pH 8), loaded onto a 5 mL Bio-Rad Mini-Proafinity IMAC column (Bio-Rad), washed with 40 mL of Buffer A-IMAC, and eluted over a 40 mL linear gradient of Buffer B-IMAC (50 mM Tris, 300 mM NaCl, 500 mM imidazole pH 8). Pure fractions were dialyzed into 20 mM HEPES, 150 mM NaCl pH 7.8.

### Protein Purification of Sso7d & nanobody mutants

Sso7d.TOM22-1, Sso7d.TOM22-2, Sso7d.c-Kit, and Sso7d.cIgY were amplified by PCR from the isolated library plasmids with forward primer Pf9 and reverse primer Pr9 and inserted into pet22B™ between the NdeI and XhoI restriction sites. NB.TOM22-1, NB.TOM22-2, NB.c-Kit-1, and NB.c-Kit-2 were amplified from isolated library plasmids by PCR with forward primer Pf10 and reverse primer Pr10 and similarly cloned into pet22B™. Positive clones were transformed into Rosetta™ *E. coli* cells for expression. Expression was carried out in 2XYT media (10 g/L tryptone, 10 g/L yeast extract, 5 g/L NaCl) plus 1X ampicillin. A 5 mL overnight culture was used to inoculate a 1 L culture of 2XYT. The cells were induced using 0.5 mM IPTG when the OD_600_ reached between 0.6 and 0.8. Expression took place overnight at 20°C. The Sso7d proteins were purified by cation exchange and metal affinity chromatography while the nanobody proteins were only purified by metal affinity chromatography.

Induced Sso7d cell culture was collected and lysed using a French press into 35 mL of Buffer A-cat (50 mM Tris-HCl, 50 mM NaCl, pH 8), loaded onto a 5 mL Bio-Rad High S column (Bio-Rad), washed with 40 mL of Buffer A-cat, and eluted with a 40 mL linear gradient of Buffer B-cat (50 mM Tris-HCl, 1 M NaCl, pH 8) from 0-50% B. The fractions were analyzed by SDS-Page and fractions containing the protein of interest were pooled and loaded onto a 5 mL Mini-profinity IMAC column (Bio-Rad). Metal affinity chromatography was performed as previously described for the TEV protease. Because the nanobody proteins were only purified using IMAC, these proteins were initially lysed into 35 mL of Buffer-A-IMAC using a French press. Pure fractions were dialyzed into PBS pH 7.4. All protein purification steps were completed at room temperature.

Each protein, expect Sso7d.c-Kit, was biotinylated using a 5:1 molar excess of Ez-Link™ Sulfo-NHS-LC-Biotin (Thermo-Fisher Scientific) overnight at 4°C. Excess reagent was inactivated through dialysis against 50 mM Tris-HCl, 300 mM NaCl pH 7.5. The purified protein was concentrated using a Vivaspin 6, 3 kDa MWCO concentrator (Sartorius). Protein concentrations were estimated by BCA assay.

### Estimation of K_D_

K_D_ values of the isolated TOM22 Sso7d, TOM22 nanobody, and c-Kit nanobody mutants were estimated using yeast surface titrations. Depending on the binder type, pCT302-TOM22 or pCT302-c-Kit yeast cells were labeled with varying concentrations of the biotinylated Sso7d or nanobody mutants followed by secondary labeling with a 1:250 dilution of streptavidin R-phycoerythrin conjugate (SA-PE; Thermo Fisher Scientific). Flow cytometry analysis was completed as previously described^31^. All labeling was performed in 50 μL of 0.1% PBSA. At low labeling concentrations (≤10 nM), yeast displaying the target of interest were combined with un-induced yeast cells for ease of obtaining cell pellets by centrifugation. Under these conditions, the volume of the labeling reaction was sufficient to ensure that the Sso7d binder mutants were present in at least 10-fold excess over the cell surface targets. Samples were incubated at room temperature for 1 hour. The K_D_ of the binding interaction between the binder protein and its target was estimated using the following relationship:

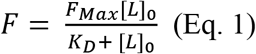

Where *F* is background subtracted mean observed fluorescence, *[L]*_0_ is the concentration of Sso7d mutant used for labeling the cells, and *F_Max_* is the background subtracted fluorescence intensity when surface saturation is attained. The fluorescence values were normalized to the highest observed fluorescence value in each data set. A global non-linear least squares regression was used to fit the data to Eq. 1 as previously described^31^. A single K_D_ value and F_max_ corresponding to each titration were used as the fitted parameters.

A complete yeast titration was not generated for Sso7d.c-Kit due to its low affinity. 2×10^6^ pCT302-c-Kit yeast cells were incubated with 2, 5, 7.5, 10, and 15 μM of Sso7d.c-Kit. Sso7d.c-Kit was not biotinylated as one of its mutated residues was a lysine. Binding of Sso7d.c-Kit to the c-Kit displaying cells was detected using an anti-penta-His Alexa 647 antibody and flow cytometry analysis as previously described.

Similar yeast surface titration analysis was also used to evaluate binding of Sso7d.cIgY to yeast cells displaying TOM22 and c-Kit. Concentrations of Sso7d.cIgY tested ranged from 4 nM to 2 μM. Because Sso7d.cIgY was biotinylated, binding was detected using SA-PE.

### Immunoblotting analysis of mitochondria isolation

2 μL of streptavidin Hi-Sur Mag 0.15 μm beads (Ocean Nanotech) were washed two times with 0.1% PBSA and incubated overnight with 80 nM of Sso7d.TOM22-1, Sso7d.TOM22-2, NB.TOM22-1, or NB.TOM22-2 diluted in 100 μL using 0.1% PBSA. Thereafter, the beads were washed two times with 0.1% PBSA and blocked with 1 mL of 1% PBSA for two hours. Subsequently, the beads were washed two times with 0.1% PBSA, resuspended in 1 mL of 0.1% PBSA, and incubated with the cell lysate. As a control, 2 μL of plain streptavidin magnetic beads (i.e. without binder functionalization) were also incubated overnight in 100 μL of 0.1% PBSA and also subjected to similar washing and blocking prior to incubation with the cell lysate. As a benchmark control, 25 μL of anti-TOM22 magnetic microbeads (Miltenyi Biotec) were incubated with the cell lysate per the manufacturers recommended protocols^49^. For each experimental condition, two bead samples were generated and incubated separately with cell lysate to allow separate analysis of mitochondrial and endoplasmic reticulum binding when immunoblotting.

To generate the cell lysate, HEK-293T cells were grown to 70-90% confluency in Dulbecco’s modified Eagle medium containing 10% fetal bovine serum (Thermo Fisher Scientific), washed with PBS, and lifted from the plate using Gibco TrypLe Select (Thermo Fisher Scientific). The cells were washed twice with ice cold PBS and resuspended in ice cold PBS containing cOmplete, mini protease inhibitor cocktail (Roche) at a concentration of 1×10^7^ cells/mL. 1 mL of cells was sheared through a 26 G needle at a time on ice about 20 times. A small aliquot of lysed cells was taken to ensure that ~80% of the cells were lysed.

1 mL of crude lysate was mixed with 8 mL of 0.1% PBSA and incubated with either the Sso7d immobilized magnetic beads, nanobody immobilized magnetic beads, or plain magnetic beads. For the commercially available TOM22 microbeads, 1 mL of crude lysate was mixed with 8.975 mL of 0.5% PBSA with 2 mM EDTA pH 7.2 (PEB buffer), per the manufacturer’s recommendations. Incubations took place for 1 hour at 4 °C with rotation. After, the streptavidin Hi-Sur bead samples were centrifuged at 13,000 g for 1 minute and washed 3x with 0.1% PBSA. The anti-TOM22 microbead samples were recovered using the manufacturer’s recommendations, supplied columns, and supplied magnet (Miltenyi Biotec). The column was placed into the magnet and washed with 3X with 1 mL of PEB buffer. The sample was then loaded onto the column adding 3.3 mL at a time and washed 3X with 3.3 mL of PEB buffer. Subsequently, the column was removed from the magnet and 1.5 mL of PEB buffer was pushed through the column using the provided syringe to release the beads into a collection tube.

After these initial washes, all samples, including the binder functionalized streptavidin beads and the anti-TOM22 microbeads, were pelleted twice at 13,000 g for 1 minute and washed twice with 0.32 M sucrose, 1 mM EDTA, and 10 mM Tris-HCl. The samples were pelleted again and resuspended in 9.75 μL of PBS and 3.75 μL of NuPAGE LDS sample buffer (Thermo Fisher Scientific) prior to boiling for 15 minutes at 98°C. The samples were cooled at room temperature for 10 minutes and then 1.5 μL of 50 mM TCEP was added. After a 10-minute incubation, the samples were loaded onto a 4-12% Bis Tris NuPAGE gel (Thermo Fisher Scientific) along with 10 μL of Novex™ Sharp Pre-stained Protein Standard (Thermo Fisher Scientific).

Two gels were run with each having lanes corresponding to the following experimental conditions: ladder, commercial anti-TOM22 micro beads, NB.TOM22-1, NB.TOM22-2, Sso7d.TOM22-1, Sso7d.TOM22-2, and plain streptavidin magnetic beads. Gels were blotted onto a PDVF membrane using the iBlot system’s program 3 for 6.5 minutes. After blotting, the membranes were blocked in 5% milk TBST (50 mM Tris HCl, 150 mM NaCl, 0.05% Tween 20 pH 7.4) for three hours. One membrane was incubated with a 1:200 dilution of mouse anti-TOM22 HRP (Santa Cruz Biotechnology; Dallas, Texas; Catalog number sc-58308 HRP; Lot number F2018) while the other membrane was incubated with a 1:200 dilution of mouse anti-calnexin HRP (Santa Cruz Biotechnology; Catalog number sc-23954 HRP; Lot number E0218) both in 15 mL of 2% milk TBST overnight. The next morning the membranes were flipped and allowed to incubate for three hours before being washed 3 times with PBS for 10 minutes each. Detection was carried out using the SuperSignal™ West Femto maximum sensitivity substrate (Thermo Fisher Scientific) according to the manufacturer’s protocol. The West-Femto substrate was left on the membranes for 1 minute prior to imaging using a Syngene Gbox with an exposure time of 2 milliseconds and the West Femto protocol. Thereafter, the gel was illuminated with white light to image the ladder only.

For the images presented in this text, the blot images were inverted. The ladder images were aligned with the blot images, the regions of interest were clipped, and irrelevant lanes were removed. No other manipulation of the images occurred.

## Supporting information

Supporting information associated with this manuscript includes:

Supplementary Methods

Supplementary Tables (2)

Supplementary Figures (4)

## Acknowledgments

This work was funded by a grant from the National Science Foundation (CBET 1511227). KBB kindly acknowledges support from an NSF Graduate Research Fellowship and a National Institute of Health Molecular Biotechnology Training Fellowship (NIH T32 GM008776).

