## Supplementary Information for "Screening yeast display libraries against magnetized yeast cell targets enables efficient isolation of membrane protein binders"

### SUPPORTING INFORMATION

**Contents:**

Supplementary Methods and Discussion

4 Supplementary Figures

2 Supplementary Tables

### Supplementary Methods and Discussion

*Evaluation of magnetization over time*

$1x10^7$ pCT302-TOM22 yeast were magnetized in 2 mL of 50 mM sodium acetate, 50 mM NaCl pH 5 buffer by incubating 100 µL of a 4 mg/mL iron oxide solution for 30 minutes at room temperature. $1x10^7$ pCT302-SsoFe2-T2A-TOM22 yeast cells were similarly magnetized, but using 50 mM Tris-HCl, 300 mM NaCl pH 7.4 (TN buffer). Two separate samples were magnetized per cell type. Cells complexed with iron oxide were removed from the unbound cells using a magnet and were washed with the appropriate magnetization buffer. After a single wash, one sample was serially diluted and plated onto SD(-Leu) plates to quantify the initial number of cells magnetized. The other sample was carried forward and washed two further times to simulate the wash and incubation steps that would occur during a typical library selection. After, this sample was incubated with 1% TNBSA pH 7.4. (TN buffer + 1% BSA) for one hour at room temperature. Next, this sample was washed 2X with 0.1% TNBSA pH 7.4 (TN buffer + 0.1% BSA) and incubated for 2 hours at room temperature in this buffer. This sample was finally washed three more times with 0.1% TNBSA, resuspended in 1 mL 0.1% TNBSA, serially diluted, and plated onto SD (-Leu) plates to quantify the cells that remained bound to the iron oxide.

*Pull downs using magnetic streptavidin beads functionalized with c-Kit nanobodies*

The c-Kit nanobodies were tested for their ability to isolate yeast cells displaying c-Kit from a heterogeneous population. Briefly, 25 µL of biotin binder Dynabeads (Thermo Fisher Scientific) were washed two times with 0.1% PBSA (PBS pH 7.4, 0.1% BSA). The beads were functionalized with either 1 µM NB.c-Kit-1, NB.c-Kit-2, or NB.TOM22-1 diluted in 100 µL of 0.1% PBSA overnight at 4°C. The beads were washed 2X with 0.1% PBSA the next day. $1x10^5$ c-Kit displaying yeast were mixed with $2x10^6$ non-displaying EBY100 yeast in 1 mL of 0.1% PBSA and added to the functionalized binder beads for 1 hour at room temperature. After one hour, the tube was placed on a magnet and the supernatant was removed. The supernatant was transferred to a new set of functionalized binder beads and a second incubation was carried out. In all, the cell mixture was incubated with three different sets of beads all functionalized with the same binding protein.

After the third incubation, the supernatant containing the cells that did not bind to the functionalized beads was centrifuged, washed 1X with 0.1% PBSA, and then labeled with a 1:100 dilution of chicken anti-c-myc antibody for 15 minutes at room temperature. Subsequently, the cells were washed and secondary labeling was carried out with a 1:250 dilution of goat-anti-chicken 488 for 10 minutes on ice. All labeling was conducted in 50 µL of 0.1% PBSA. Anti-c-myc labeling was also performed on a cell population that did not undergo incubation with the binder proteins that contained $1x10^5$ c-Kit displaying yeast and $2x10^6$ EBY100 yeast. The EBY100 cells do not express c-myc, so any c-myc expression is attributed to cells expressing c-Kit. If c-Kit expressing cells were pulled out by the functionalized magnetic beads then the percentage of c-myc expressing cells would be greater in the initial population when compared to the population of cells that were not pulled out by the beads. Thus, the percent decrease in cells expressing c-myc between the final and initial populations was quantified to assess the depletion of c-Kit displaying yeast cells by the binder functionalized beads.

The c-Kit cell population was modestly reduced through incubation with magnetic beads functionalized with the c-Kit specific nanobodies. Specifically, there was a ~ 10% reduction in c-Kit displaying cells after incubation with magnetic beads functionalized with Nb.c-KIT.1 and NB-c-Kit.2, respectively, at room temperature (**Figure S3**). However, there was no statistically significant difference in the recovery of c-Kit displaying cells by magnetic beads functionalized with the c-Kit specific nanobodies in comparison to a TOM22 specific nanobody. The nanobodies were also immobilized onto His dynabeads (Thermo Fisher Scientific) following a similar procedure (2 μL His-Tag Isolation Dynabeads, 2.3 μM nanobody protein) to understand if immobilization onto streptavidin beads was impeding the nanobody binding (**Figure S4**). NB.TOM22-2 was used as the negative control and the incubations were carried out cold. The cell populations were only incubated twice with the functionalized beads, instead of three incubations. However, the decrease in c-myc expression between the final and initial populations was similar when the nanobodies were functionalized onto His dynabeads and streptavidin dynabeads. In general, the functionalization method had no influence on the recovery of the c-KIT displaying cells. While these combined results suggest some specificity of the nanobody mutants for c-Kit, it is likely that higher affinity ligands are required for significant enrichment of c-Kit displaying cells from heterogenous populations. Hence, further evaluation of the c-Kit nanobodies was not carried out to characterize their ability to isolate mammalian cells that naturally express c-Kit.

### Supplementary Figures

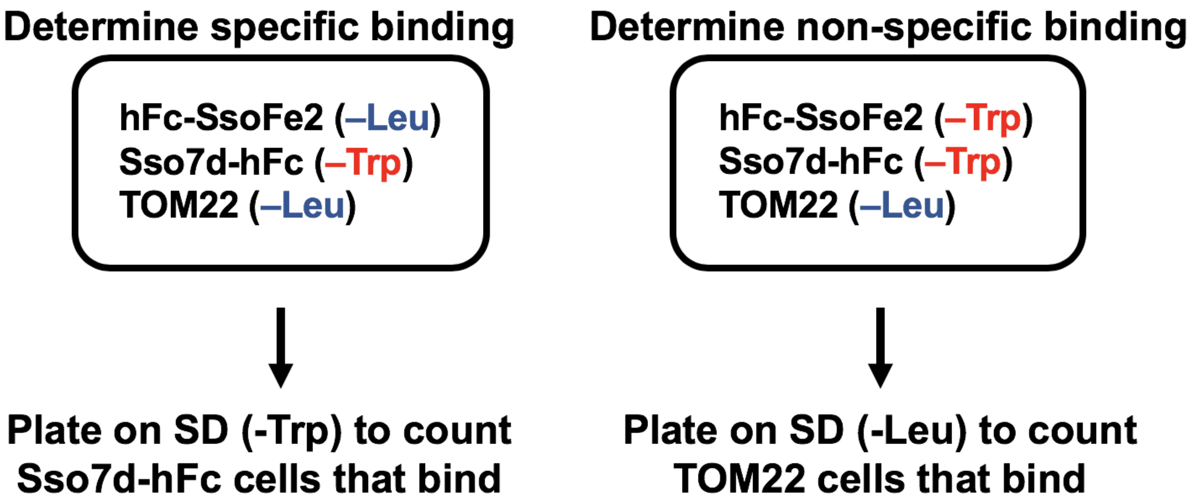


**Figure S1**. Experimental conditions and plating methods used to quantify specific (Sso7d-hFc displaying yeast) and non-specific binding (TOM22 displaying yeast) to magnetized yeast cells co-displaying the target protein (hFc) and SsoFe2.

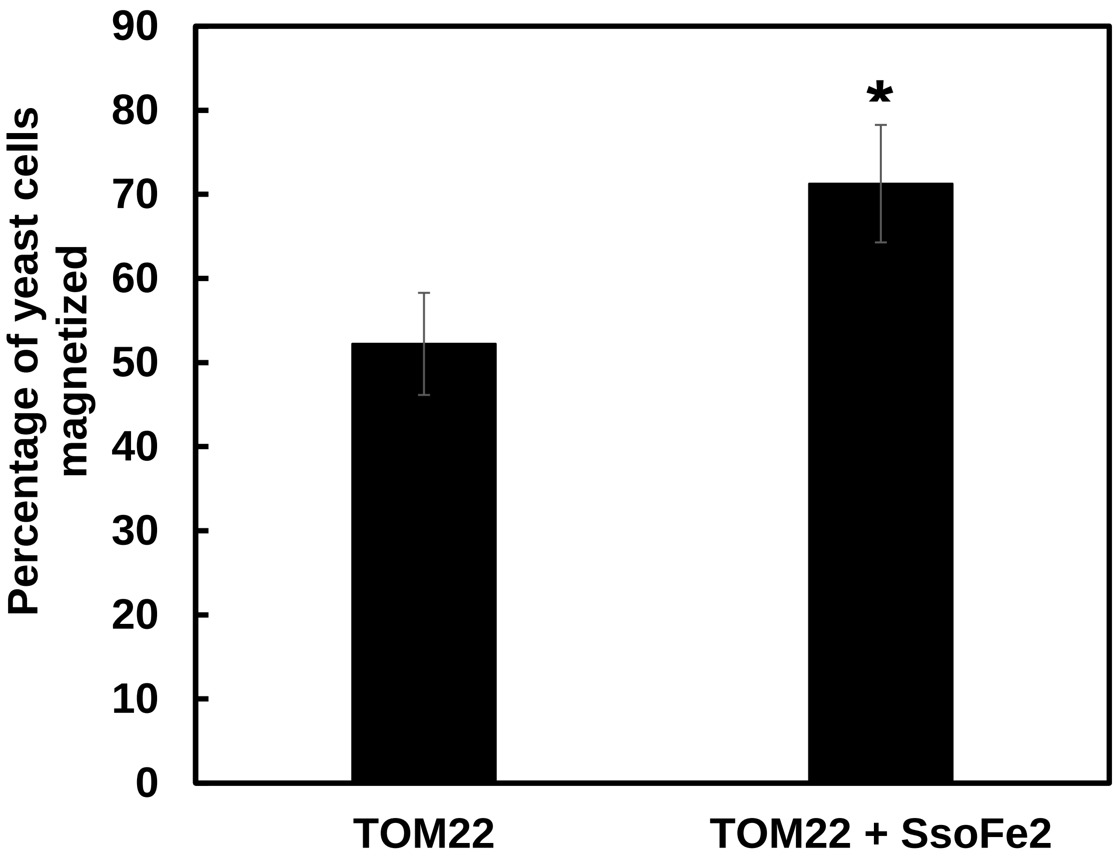


**Figure S2**. Percentage of yeast initially magnetized that remained magnetized over the course of the wash, blocking, and incubation steps that would be associated with a library screen. Yeast cells expressing only TOM22 and yeast cells expressing both TOM22 and SsoFe2 were evaluated for their magnetization efficiency. Both cell populations were magnetized after incubation with iron oxide nanoparticles. SsoFe2 affords affinity-based magnetization while the cells not displaying SsoFe2 are magnetized through electrostatic interactions. Error bars represent the standard error of the mean for six repeats for cells displaying TOM22 only and three repeats for cells displaying TOM22, SsoFe2. * represents $p < .01$ for two tailed t-test in comparison to TOM22 only population.

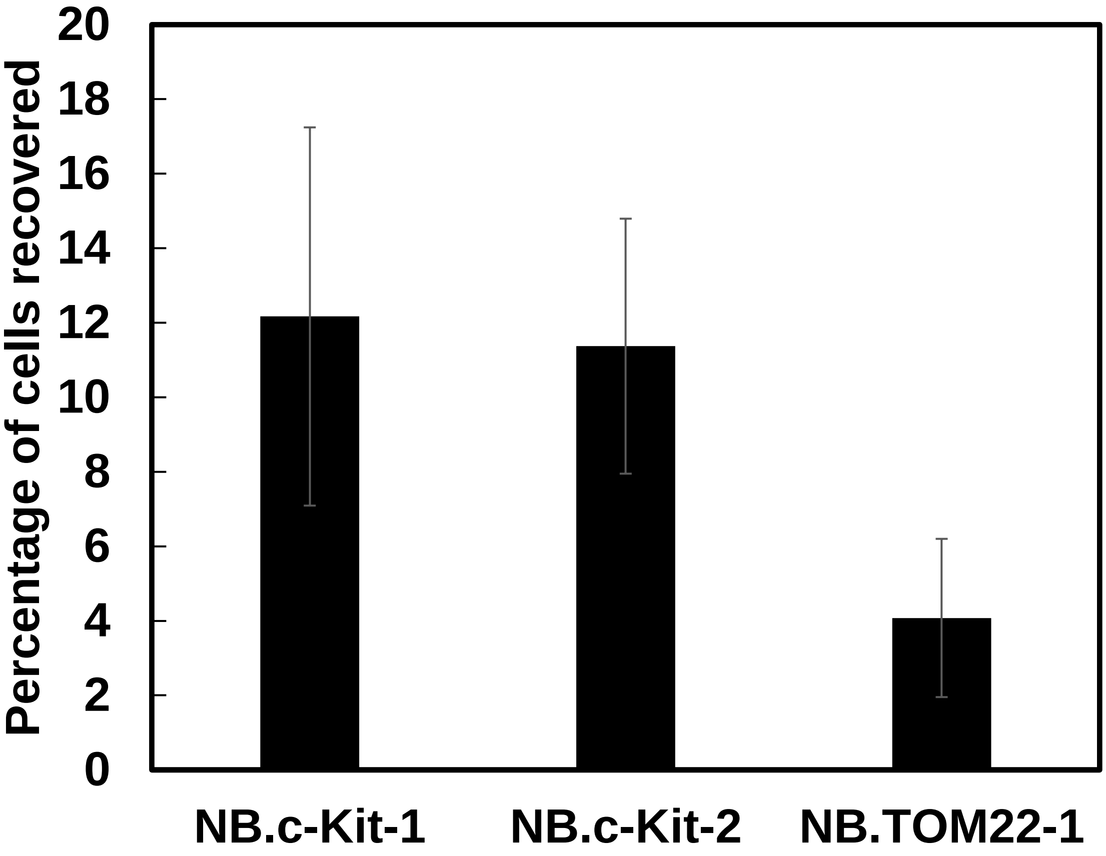


**Figure S3**. Percentage of c-Kit displaying cells recovered from a mixed population using magnetic streptavidin beads functionalized with binders NB.c-Kit-1, NB.c-Kit-2, and NB.TOM22-1. The initial population contained $1x10^5$ c-Kit displaying yeast and $2x10^6$ non-displaying EBY100 yeast. The reduction in c-Kit displaying cells was quantified by comparing the c-myc expression between the unrecovered cell population and the initial population. These results suggest that higher affinity binders may be required to obtain more significant pull out of specific cell types for heterogenous populations. Error bars represent the standard error of the mean for three repeats.

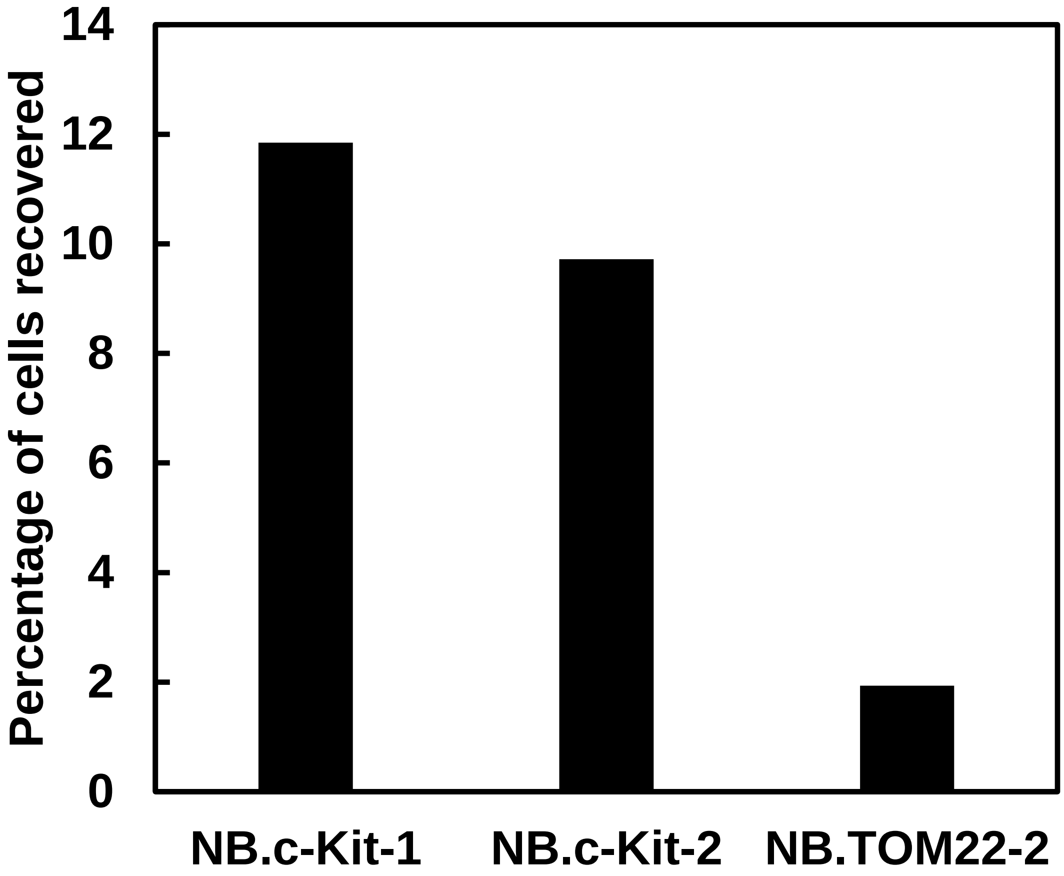


**Figure S4**. Percentage of c-Kit displaying cells recovered from a mixed population using magnetic His beads functionalized with binders NB.c-Kit-1, NB.c-Kit-2, and NB.TOM22-2. The cell populations were only incubated with two sets of functionalized beads in comparison to the previous experiments that involved three bead incubations. Data from a single experiment is shown.

### Supplementary tables

**Table S1**. List of gene fragments

| | |
|---|---|
| Gene Block 1 | GAATTCATGAAGGTTTTGATTGTCTTGTTGGCTATCTTCGCTGCTTT<br>GCCATTGGCCTTAGCTCAACCAGTAATTTCTACTACCGTCGGTTCC<br>GCTGCAGAAGGCTCTTTGGACAAGAGACAGGAACTGACAACTATA<br>TGCGAGCAAATCCCCTCACCAACTTTAGAATCGACGCCGTACTCTT<br>TGTCAACGACTACTATTTTGGCCAACGGGAAGGCAATGCAAGGAG<br>TTTTTGAATATTACAAATCAGTAACGTTTGTCAGTAATTGCGGTTC<br>TCACCCCTCAACAACTAGCAAAGGCAGCCCCATAAACACACAGTA<br>TGTTTTTTATCCGTACGACGTTCCAGACTACGCTGGTGGTGGTGGT<br>TCCGGTGGTGGTGGTTCTGGTGGTGGTGGTTCAGCTAGCGCTGCCG<br>CCGTCGCTGCTGCCGGTGCAGGGGAACCCCAGTCCCCGGACGAAT<br>TGCTCCCGAAAGGCGACGCGGAGAAGCCTGAGGAGGAGCTGGAG<br>GAGGACGACGATGAGGAGCTAGATGAGACCCTGTCGGAGAGACT<br>ATGGGGCCTGACGGAGATGTTTCCGGAGAGGGTCCGGTCCGCGGC<br>CGGAGCCACTTTTGATCTTTCCCTCTTTGTGGCTCAGAAAATGTAC<br>AGGTTTTCCAGGGCAGGATCCGAACAAAAGCTTATCTCCGAAGAA<br>GACTTGGAGGGAAGGGGATCTTTGCTAACGTGCGGGGATGTCGAA<br>GAAAACCCTGGCCCTATGAAGGTGCTGATTGTTCTATTGGCTATAT<br>TTGCTGCCTTGCCCCTGGCCCTGGCACAGCCAGTCATAAGTACAAC<br>AGTGGGCAGTGCTGCGGAAGGTTCACTAGATAAAAGACAGGAACT<br>TACTACAATTTGTGAACAAATACCTAGTCCCACGTTGGAGTCAACA<br>CCGTACTCCCTGTCTACTACAACAATACTTGCGAACGGTAAGGCCA<br>TGCAAGGAGTCTTTGAATACTACAAGTCAGTTACCTTCGTATCTAA<br>CTGCGGGAGTCACCCGAGCACCACATCCAAGGGGAGTCCTATAAA<br>CACTCAGTATGTATTCGACTATAAGGATGACGATGACAAGGGCGG<br>CGGAGGCTCCGGTGGAGGCGGAAGCGGCGGTGGAGGTAGTCCTA<br>GGGCGACCGTGAAATTTAAATATAAAGGCGAAGAAAAACAGGTG<br>GATATTAGCAAAATTAGACGGGTGCGTCGCAAGGGCAAATGCATT<br>AGGTTTTACTATGATCTGGGCGGCGGCAAATATGGCAGGGGCATA<br>GTGAGCGAAAAAGATGCGCCGAAAGAACTGCTGCAGATGCTGGA<br>AAAACAGAAAAAACATATGGGTAAACCCATACCCAATCCCCTGCT<br>GGGTCTTGATAGTACGTAATAG |
| Gene Block 2 | CCCAAATCTTGTGACAAAACTCACACATGCCCACCGTGCCCAGCA<br>CCTGAACTCCTGGGGGGACCGTCAGTCTTCCTCTTCCCCCCAAAAC<br>CCAAGGACACCCTCATGATCTCCCGGACCCCTGAGGTCACATGCG<br>TGGTGGTGGACGTGAGCCACGAAGACCCTGAGGTCAAGTTCAACT<br>GGTACGTGGACGGCGTGGAGGTGCATAATGCCAAGACAAAGCCGC<br>GGGAGGAGCAGTACAACAGCACGTACCGTGTGGTCAGCGTCCTCA<br>CCGTCCTGCACCAGGACTGGCTGAATGGCAAGGAGTACAAGTGCA<br>AGGTCTCCAACAAAGCCCTCCCAGCCCCCATCGAGAAAACCATCT<br>CCAAAGCCAAAGGGCAGCCCCGAGAACCACAGGTGTACACCCTGC<br>CCCCATCCCGGGATGAGCTGACCAAGAACCAGGTCAGCCTGACCT<br>GCCTGGTCAAAGGCTTCTATCCCAGCGACATCGCCGTGGAGTGGG<br>AGAGCAATGGGCAGCCGGAGAACAACTACAAGACCACGCCTCCC |

| | |
|---|---|
| | GTGCTGGACTCCGACGGCTCCTTCTTCCTCTACAGCAAGCTCACCG<br>TGGACAAGAGCAGGTGGCAGCAGGGGAACGTCTTCTCATGCTCCG<br>TGATGCATGAGGCTCTGCACAACCACTACACGCAGAAGAGCCTCT<br>CCCTGTCTCCGGGTAAA |
| Gene Block 3 | CAACCATCTGTGAGTCCAGGGGAACCGTCTCCACCATCCATCCATC<br>CAGGAAAATCAGACTTAATAGTCCGCGTGGGCGACGAGATTAGGC<br>TGTTATGCACTGATCCGGGCTTTGTCAAATGGACTTTTGAGATCCT<br>GGATGAAACGAATGAGAATAAGCAGAATGAATGGATCACGGAAA<br>AGGCAGAAGCCACCAACACCGGCAAATACACGTGCACCAACAAA<br>CACGGCTTAAGCAATTCCATTTATGTGTTTGTTAGAGATCCTGCCA<br>AGCTTTTCCTTGTTGACCGCTCCTTGTATGGAAAGAAGACAACGA<br>CACGCTGGTCCGCTGTCCTCTCACAGACCCAGAAGTGACCAATTAT<br>TCCCTCAAGGGGTGCCAGGGGAAGCCTCTTCCCAAGGACTTGAGG<br>TTTATTCCTGACCCCAAGGCGGGCATCATGATCAAAAGTGTGAAA<br>CGCGCCTACCATCGGCTCTGTCTGCATTGTTCTGTGGACCAGGAGG<br>GCAAGTCAGTGCTGTCGGAAAAATTCATCCTGAAAGTGAGGCCAG<br>CCTTCAAAGCTGTGCCTGTTGTGTCTGTGTCCAAAGCAAGCTATCT<br>TCTTAGGGAAGGGGAAGAATTCACAGTGACGTGCACAATAAAAGA<br>TGTGTCTAGTTCTGTGTACTCAACGTGGAAAAGAGAAAACAGTCA<br>GACTAAACTACAGGAGAAATATAATAGCTGGCATCACGGTGACTT<br>CAATTATGAACGTCAGGCAACGTTGACTATCAGTTCAGCGAGAGT<br>TAATGATTCTGGAGTGTTCATGTGTTATGCCAATAATACTTTTGGA<br>TCAGCAAATGTCACAACAACCTTGGAAGTAGTAGATAAAGGATTC<br>ATTAATATCTTCCCCATGATAAACACTACAGTATTTGTAAACGATG<br>GAGAAAATGTAGATTTGATTGTTGAATATGAAGCATTCCCCAAAC<br>CTGAACACCAGCAGTGGATCTATATGAACAGAACCTTCACTGATA<br>AATGGGAAGATTATCCCAAGTCTGAGAATGAAAGTAATATCAGAT<br>ACGTAAGTGAACTTCATCTAACGAGATTAAAAGGCACCGAAGGAG<br>GCACTTACACATTCCTAGTGTCCAATTCTGACGTCAATGCTGCCAT<br>AGCATTTAATGTTTATGTGAATACAAAACCAGAAATCCTGACTTAC<br>GACAGGCTCGTGAATGGCATGCTCCAATGTGTGGCAGCAGGATTC<br>CCAGAGCCCACAATAGATTGGTATTTTTGTCCAGGAACTGAGCAG<br>AGATGCTCTGCTTCTGTACTGCCAGTGGATGTGCAGACACTAAACT<br>CATCTGGGCCACCGTTTGGAAAGCTAGTGGTTCAGAGTTCTATAGA<br>TTCTAGTGCATTCAAGCACAATGGCACGGTTGAATGTAAGGCTTAC<br>AACGATGTGGGCAAGACTTCTGCCTATTTTAACTTTGCATTTAAAG<br>GTAACAACAAAGAGCAAATCCATCCCCACACCCTGTTCACTCCT |
| Gene Block 4 | ATGAAGGTTTTGATTGTCTTGTTGGCTATCTTCGCTGCTTTGCCATT<br>GGCCTTAGCTCAACCAGTAATTTCTACTACCGTCGGTTCCGCTGCA<br>GAAGGCTCTTTGGACAAGAGACAGGAACTGACAACTATATGCGAG<br>CAAATCCCCTCACCAACTTTAGAATCGACGCCGTACTCTTTGTCAA<br>CGACTACTATTTTGGCCAACGGGAAGGCAATGCAAGGAGTTTTTG<br>AATATTACAAATCAGTAACGTTTGTCAGTAATTGCGGTTCTCACCC<br>CTCAACAACTAGCAAAGGCAGCCCCATAAACACACAGTATGTTTT<br>TCCCGGGTATCCGTACGACGTTCCAGACTACGCTGGTGGTGGTGGT<br>TCCGGTGGTGGTGGTTCTGGTGGTGGTGGTTCAGCTAGCGCTGCCG |

| | |
|---|---|
| | CCGTCGCTGCTGCCGGTGCAGGGGAACCCCAGTCCCCGGACGAATTGCTCCCGAAAGGCGACGCGGAGAAGCCTGAGGAGGAGCTGGAGGAGGACGACGATGAGGAGCTAGATGAGACCCTGTCGGAGAGACTATGGGGCCTGACGGAGATGTTTCCGGAGAGGGTCCGGTCCGCGGCCGGAGCCACTTTTGATCTTTCCCTCTTTGTGGCTCAGAAAATGTACAGGTTTTCCAGGGCAGGATCCGAACAAAAGCTTATCTCCGAAGAAGACTTGTAATGACTCGAG |
| Gene Block 5 | CTATTACGTACTATCAAGACCCAGCAGGGGATTGGGTATGGGTTTACCCCATGGTTTTTTCTGTTTTTCCAGCATCTGCAGCAGTTCTTTCGGCGCATCTTTTTCGCTCACTATGCCCCTGCCATATTTGCCGCCGCCCAGATCATAGTAAAACCTAATGCATTTGCCCTTGCGACGCACCCGTCTAATTTTGCTAATATCCACCTGTTTTTCTTCGCCTTTATATTTAAATTTCACGGTCGCCCTAGGACTACCTCCACCGCCGCTTCCGCCTCCACCGGAGCCTCCGCCGCCCTTGTCATCGTCATCCTTATAGTCGAATACATACTGAGTGTTTATAGGACTCCCCTTGGATGTGGTGCTCGGGTGACTCCCGCAGTTAGATACGAAGGTAACTGACTTGTAGTATTCAAAGACTCCTTGCATGGCCTTACCGTTCGCAAGTATTGTTGTAGTAGACAGGGAGTACGGTGTTGACTCCAACGTGGGACTAGGTATTTGTTCACAAATTGTAGTAAGTTCCTGTCTTTTATCTAGTGAACCTTCCGCAGCACTGCCCACTGTTGTACTTATGACTGGCTGTGCCAGGGCCAGGGGCAAGGCAGCAAATATAGCCAATAGAACAATCAGCACCTTCATAATTCCTTGGAATTTTCAAAAATTCTTACTTTTTTTTTTGGATGGACGCAAAGAAGTTTAATAATCATATTACATGGCATTACCACCATATACATATCCATATACATATCCATATCTAATCTTACTTATATGTTGTGGAAATGTAAAGAGCCCCATTATCTTAGCCTAAAAAAACCTTCTCTTTGGAACTTTCAGTAATACGCTTAACTGCTCATTGCTATATTGAAGTACGGATTAGAAGCCGCCGAGCGGGTGACAGCCCTCCGAAGGAAGACTCTCCTCCGTGCGTCCTCGTCTTC |

**Table S2**. List of oligonucleotide primers

| Primer | Sequence |
|---|---|
| Pf1 | TCTTATTCAAATGTAATAAAAGATCGAATTCATGAAGGTTTTGATTGTC |
| Pr1 | TACATCTACACTGTTGTTATCAGATCTCGAGCTATTACGTACTATCAAGACCC |
| Pf2 | GTTGACGCTAGCCCCAAATCTTGTGACAAAACTCAC |
| Pr2 | GCACTTGGATCCTTTACCCGGAGACAGGGAGA |
| Pf3 | GTTGACGCTAGCCAACCATCTGTGAGTCCAGGG |
| Pr3 | GCACTTGGATCCAGGAGTGAACAGGGTGTGGG |
| Pf4 | GCTACGGAATTCATGAAGGTTTTGATTGTCTTGTTGGC |
| Pr4 | GCTACGCTCGAGTCATTACAAGTCTTCTTCGGAGATAA |
| Pf5 | GCTATCGGTACCCTATTACGTACTATCAAGACCCAG |
| Pr5 | GTACTGACCGGTGAAGACGAGGACGCACGGA |
| Pf6 | GAGTCAGCTAGCATGGCGACCGTGAAATTTAAATAT |
| Pr6 | GAGTCAGGATCCTTTTTTCTGTTTTTCCAGCATCTG |

| | |
|---|---|
| Pf7 | GTTCTCGCTAGCGCTGCCGCCGTCGCTG |
| Pr7 | GCACTTGGATCCTGCCCTGGAAAACCTGTACATTT |
| Pf8 | CAGAAGGCTCTTTGGACAAGAGACACCACCACCACCACCACGCTAGCAT GGCGACCGTGAAATTTAAATAT |
| Pr8 | GGAGATAAGCTTTTGTTCCCCTTGGAAGTACAGGTTCTCGGATCCTTTTT TCTGTTTTTCCAGCATCTG |
| Pf9 | GATGCACATATGATGGCGACCGTGAAATTT |
| Pr9 | CCGCCGCTCGAGTTTTTTCTGTTTTTCCAGCATC |
| Pf10 | GTCTCGCATATGGCACAAGTTCAGCTTGTAGAGT |
| Pr10 | GTCACTCTCGAGCGATGATACAGTTACTTGGGTAC |
